# Ensemble-based genomic prediction for maize flowering-time improves prediction accuracy and reveals novel insights into trait genetic variation

**DOI:** 10.1101/2025.07.15.664852

**Authors:** Shunichiro Tomura, Owen Powell, Melanie J. Wilkinson, Mark Cooper

## Abstract

While various genomic prediction models have been evaluated for their potential to accelerate genetic gain for multiple traits, no individual genomic prediction model has outperformed all others across all applications. As an alternative approach, ensembles of multiple individual genomic prediction models can be applied to utilise the complementary strengths of individual prediction models and offset the prediction errors of each. We used the EasiGP (Ensemble AnalySis with Interpretable Genomic Prediction) pipeline to investigate the performance of an ensemble approach, targeting flowering-time traits measured in two maize nested association mapping datasets. For both datasets, the ensemble-based prediction approach achieved higher prediction accuracy and lower prediction error across the flowering-time traits compared to each individual model. Multiple genomic regions known to contain key flowering-time related genes were repeatedly included as features across individual genomic prediction models, indicating the models successfully captured SNPs as features that are associated with genomic regions known to contain flowering-time genes. Although repeatability was high for some genomic regions, estimated marker effects varied across many genomic regions, suggesting that the models might also have captured different aspects of the genetic variation underlying the traits. The ensemble combination of the diverse views likely contributed to the improvement of prediction performance by the ensemble-based approach over the individual prediction models. Ensemble-based prediction can be applied to overcome limitations observed in the continuous exploration for the best individual genomic prediction models that can consistently achieve the highest prediction performance, thereby potentially contributing to improved prediction accuracy for applications in crop breeding.

**Article summary:** This study targets researchers interested in the performance of genomic prediction models. To demonstrate potential advantages of an ensemble of diverse individual genomic prediction models, we investigated the prediction of key flowering-time traits (days to anthesis and anthesis to silking interval) in two maize datasets. The ensemble approach consistently improved the prediction performance. The improvement was attributed to the offset of prediction errors by combining multiple different dimensions of trait genetic variation. Ensembles can lead to higher selection accuracy of desirable individuals for applications in crop breeding.

## Introduction

Crop breeding programs have targeted the development of more resilient genotypes for harsh biotic and abiotic stress conditions (Kholová et al. 2021; Cooper and Messina 2023; Werner et al. 2025). The pressures of climate change have amplified such stresses in agricultural environments, resulting in increasing demand for better-adapted crop genotypes (Ceccarelli et al. 2010; Langridge et al. 2021). Consequently, breeders have endeavoured to continuously select genotypes with improved combinations of the target traits to accelerate genetic gain (Messina et al. 2023; Pixley et al. 2023).

Genomic selection (Meuwissen et al. 2001; Bernardo and Yu 2007; Voss-Fels et al. 2019; Khaipho-Burch et al. 2023; Crossa et al. 2025; Escamilla et al. 2025) is a key approach to accelerate genetic gain. The prediction of trait phenotypes using genomic markers can be used to reduce the length of each breeding cycle and therefore the cost of breeding programs (Heffner et al. 2010). Genomic prediction models are trained to estimate the effect of each genomic marker for target trait prediction by capturing patterns of genetic variation associated with phenotypic differences in the reference population of the breeding program (Alemu et al. 2024). Genomic prediction models that can identify repeatable associations between marker combinations and trait variation across populations have the potential to improve the prediction performance for multiple breeding applications (Hammer et al. 2006; Cooper et al. 2014a,b; Voss-Fels et al. 2019; Diepenbrock et al. 2022).

Numerous genomic prediction models have been investigated across a wide range of applications to crop breeding programs (Cooper et al. 2014a,b; Diepenbrock et al. 2022; John et al. 2022; Crossa et al. 2025; Escamilla et al. 2025). However, none of the proposed approaches has shown superior performance for all traits and datasets. For example, Heslot et al. (2012) showed an absence of a consistent best model across eight prediction models (parametric and machine learning) applied to wheat, barley and maize datasets. Similarly, Plavšin et al. (2022) evaluated eight prediction models for quality traits in wheat, revealing that while ridge regression best linear unbiased prediction (rrBLUP) reached the overall highest performance, the differences from other prediction models were small and the most accurate genomic prediction model was dependent on the environment and the trait. Genomic prediction performance was investigated comprehensively by Washburn et al. (2025); the global genotype-by-environment (GxE) prediction competition evaluated a wide range of genomic prediction models from 128 teams for predicting maize yield traits. Their results showed that no individual genomic prediction model consistently improved prediction performance. Instead, an ensemble of parametric and machine learning models reached the overall highest prediction performance. Other empirical studies (Meher et al., 2022; Lourenço et al., 2024; Gibbs et al., 2025) that also compared the prediction performance of individual genomic prediction models consistently found that no prediction model was superior to others across different prediction scenarios created by combinations of environments, crops and datasets. These observed results demonstrate the limitations in the development of a single genomic prediction model that is stably generalised with high prediction performance in new environments under cross-validation settings.

The lack of a best individual genomic prediction model could be a consequence of the No Free Lunch Theorem (Wolpert and Macready 1997), which states that the mean performance of prediction models becomes equivalent across all scenarios. The lack of a consistently superior model is likely a consequence of individual prediction models being well-suited for specific scenarios, but not for all. Hence, the continuous exploration for an all-rounder individual genomic prediction model may not be an optimal approach to improve overall prediction performance.

Alternatively, as shown in Washburn et al. (2025), multiple genomic prediction models can be combined as an ensemble to improve prediction performance rather than implementing them independently (Cooper et al. 2025; Messina et al. 2025). For instance, Nascimento et al. (2024) successfully improved genomic prediction performance for several traits, such as yield and total number of fruits, in an Arabica coffee breeding program using ensembles of diverse prediction models. The success of ensembles can be theoretically explained by the Diversity Prediction Theorem (Hong and Page 2004; Page 2007; Page 2018); the use of diverse prediction algorithms in a Many-Model ensemble lowers the prediction error more than the average prediction error of individual prediction models. Hence, the prediction error of an ensemble depends on the evaluated prediction errors of individual models included in the ensemble and the diversity of their predictions (Messina et al. 2025; Tomura et al. 2025b).

Using the framework of the Diversity Prediction Theorem, here we investigated whether an ensemble of diverse genomic prediction models improved prediction performance by predicting maize flowering-time traits in two well-documented Nested Association Mapping (NAM) examples (Buckler et al. 2009; Chen et al. 2019). The parents of these two NAM datasets represent a wide range of maize genetic diversity (Fig. 1), extending from domestication to modern inbred lines used for hybrid breeding (Hufford et al. 2012). This study was guided by two objectives. Firstly, the prediction performance of the ensemble was compared with individual genomic prediction models to evaluate the expectations of the Diversity Prediction Theorem for prediction performance improvement. Secondly, the estimates of genomic marker effects on the target traits from each genomic prediction model were compared to observe the association between the diversity in quantified marker effect values and the prediction performance of the ensemble.

**Fig. 1:**
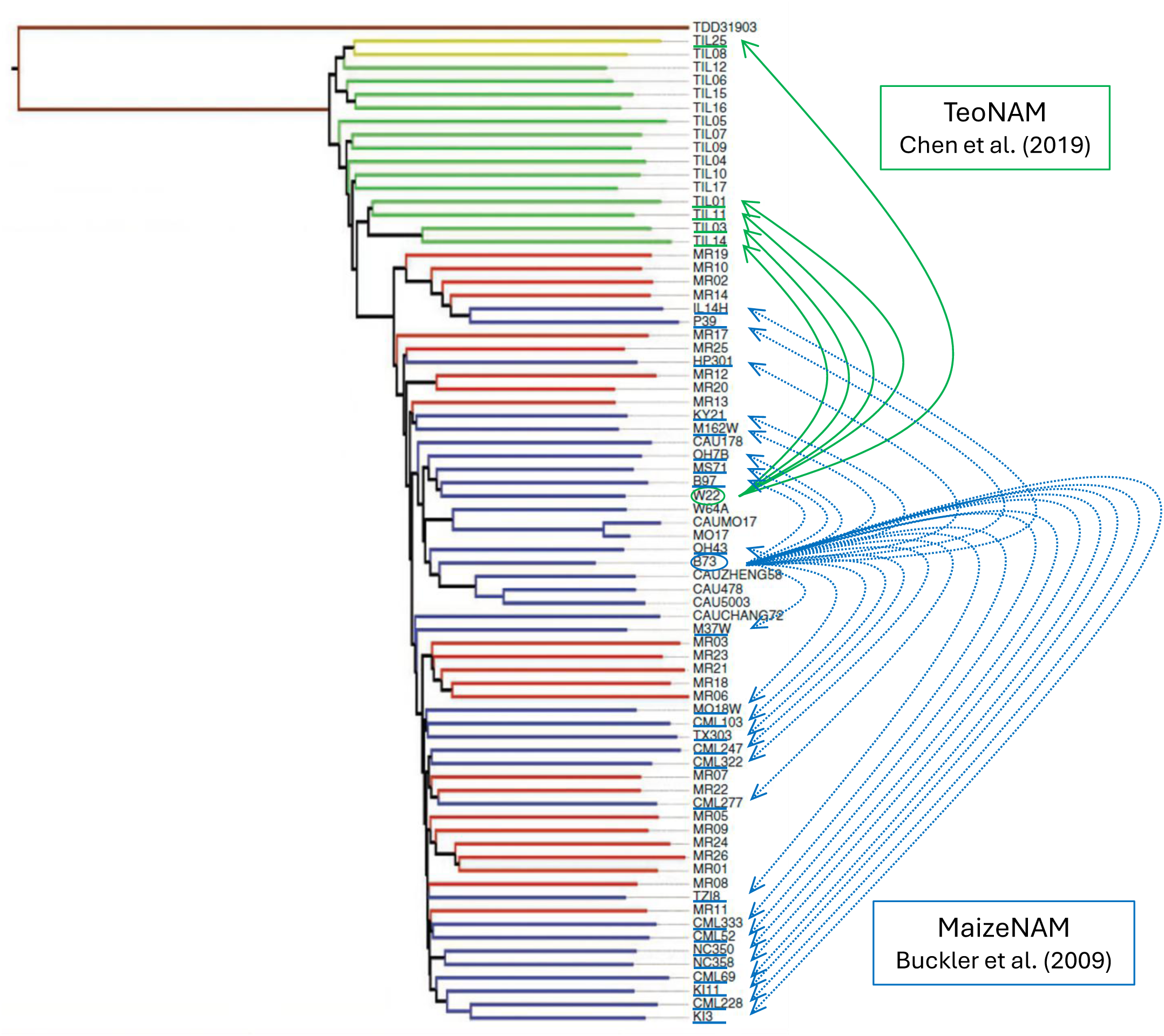
Neighbour-joining tree showing sequence relationships between TeoNAM and MaizeNAM parents (adapted from Hufford et al. (2012) with permission from Professor Jeffrey Ross-Ibarra). Within the tree, each colour represents *parviglumis* (green), landraces (red), improved lines (blue), *mexicana* (yellow) and *Tripsacum* (brown), respectively. Circles around W22 and B73 represent the common parent in the TeoNAM (Chen et al. 2019) and MaizeNAM (Buckler et al. 2009) datasets, respectively. Green and blue lines under several maize line names show donor parents for the TeoNAM and MaizeNAM datasets, respectively. For the TeoNAM dataset, W22 was crossed (green arrows) with five donor parents. For the MaizeNAM dataset, B73 was crossed (blue dash arrows) with 25 donor parents.

## Materials and Methods

### 1. Datasets

The TeoNAM dataset (Chen et al. 2019) is a collection of samples from the five recombinant inbred line (RIL) populations between the maize line W22 and five teosinte types (TIL01, TIL03, TIL11 and TIL14 from *Z. mays ssp. parviglumis* and TIL25 from *Z.mays ssp. mexicana*). In each RIL population, the F1 was backcrossed with W22. After the backcross was completed, each population was self-pollinated four times to develop the RIL populations. A randomised complete block design was applied to test the RILs for each population contributing to the TeoNAM experiment. Each population was tested twice at the University of Wisconsin West Madison Agricultural Research Station. For W22TIL01, W22TIL03 and W22TIL11, the RILs were tested in the summers of 2015 and 2016. For W22TIL14, the RILs were tested in the summers of 2016 and 2017, and for W22TIL25, the RILs were tested in two different blocks in the summer of 2017.

The MaizeNAM dataset (Buckler et al. 2009) consists of samples from 25 RIL populations derived from crosses between the maize line B73 and a diverse set of 25 inbred lines from temperate and tropical regions. The 25 RIL populations were developed through self-pollination to the F5 generation. Each population was tested in the following locations during the summer in the United States: Aurora in New York, Clayton in North Carolina, Columbia in Missouri and Urbana in Illinois. Two experiments were conducted at each location with a randomised design and hence each population was evaluated in the eight environments. The recorded phenotype scores in each environment were used to estimate the best linear unbiased predictions (BLUPs) with ASReml (v2.0) (Butler et al. 2017) in each population. The best linear unbiased estimates (BLUEs) were calculated by unshrinking the BLUPs per population using heritability, after removing the mean phenotype value for each population (Fig. S1). The combination of mean phenotype values and heritability uniquely characterised each population, showing mild positive correlations as the overall trend (Fig. S2). In the main manuscript, BLUEs were employed as the target trait phenotypes of each RIL in each prediction scenario for the MaizeNAM dataset. The comparable prediction results and analysis for the BLUPs are reported in the Supplementary Material (Fig. S3, S10, S11). The total number of genomic markers (SNPs) and RIL samples from each target dataset is summarised in Table 1 for the dataset level and Table S1 for the population level.

**Table 1:** The total number of genomic markers (SNPs) before and after linkage disequilibrium (LD) filtering and the number of recombinant inbred line (RILs) records for the TeoNAM and MaizeNAM datasets.

| Dataset | SNPs |  | RIL<br>Records |
| --- | --- | --- | --- |
|  | Original | LD Filtering |  |
| TeoNAM | 11,396 - 16,110 | 268 - 342 | 438 – 616 |
| MaizeNAM | 1,106 | 170 - 251 | 126 – 196 |

The selection of the two maize NAM datasets was motivated by the genetic diversity that covered a continuum of maize lines from wild relatives to post-domestication lines (Fig. 1; Hufford et al. 2012). The TeoNAM dataset contains the highest level of genetic diversity: Crosses of a temperate maize inbred with representatives of its ancestral species, Teosinte, can uncover components of trait genetic variation fixed during domestication. In contrast, the MaizeNAM dataset is less diverse, as crosses are limited to within a set of domesticated maize lines. Using these two maize NAM datasets, characterised by different levels of genetic diversity, we investigated the distinct characteristics of the genomic prediction models for these datasets to understand the predictive behaviour of the individual models and their ensemble.

### 2. Data preprocessing

We applied the same data preprocessing methods reported by Tomura et al. (2025b). Briefly, for the TeoNAM dataset, missing SNP markers were imputed by assigning genomic marker values based on flanking markers. RIL samples with missing genomic markers for an entire chromosome and those without phenotype values were removed. For the MaizeNAM dataset, missing SNP markers were imputed with flanking markers and missing phenotypes were removed, as reported by Buckler et al. (2009). Hence, no further imputation was required. Prior to the ensemble-based analyses, the total number of SNPs was reduced further based on linkage disequilibrium (LD) using PLINK (v1.9) (Chang et al. 2015). Any SNP with a squared correlation of 0.8 or above was removed using a window size of 30 kb and a step size of 5 SNPs in both datasets.

The two flowering-time related traits, days to anthesis (DTA) and anthesis to silking interval (ASI), were measured in both datasets. For the TeoNAM dataset, the DTA and ASI traits for each RIL were recorded across two environments as reported by Chen et al. (2019). These environments were concatenated for the ensemble genomic prediction analysis following Tomura et al. (2025b), combining genotype and phenotype data from two environments into a single dataset with a factor indicating the corresponding environment for each RIL population dataset.

### 3. Genetic variance component calculation

Both additive and non-additive (epistatic) variance components were estimated per population for both DTA and ASI in each dataset following Yadav et al. (2021, 2023). Each variance component was calculated by constructing the respective genomic relationship matrices. The additive genomic relationship matrix (𝐺_𝐴_) was defined below (Yang et al. 2010):

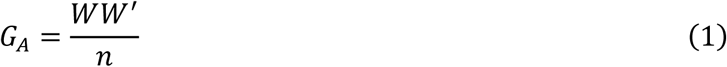

where 𝑊 is the incidence matrix corresponding to the additive marker effects and 𝑛 is the number of individuals. The epistatic genomic relationship matrix (𝐺_𝐴𝐴_) was developed by the Hadamard product (⊙) of the additive genomic relationship matrix as defined below:

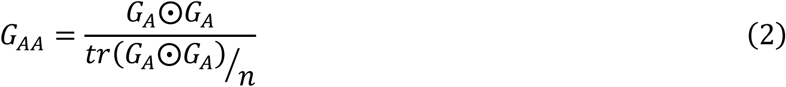

where 𝑡𝑟(·) is the trace that sums the diagonal elements in the matrix. Using the 𝐺_𝐴_ and 𝐺_𝐴𝐴_ matrices, extended genomic best linear unbiased prediction (extended-GBLUP) was modelled:

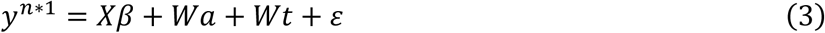

where 𝑦 is the vector of predicted phenotypes, 𝑋 is the incidence matrix assigned to fixed effects 𝛽, 𝑊 is the incidence matrix for the random effects, 𝑎∼𝑁(0, 𝐺_𝐴_, 𝜎_A_^2^) is the vector of additive genomic marker effects, 𝑡∼𝑁(0, 𝐺_𝐴𝐴_, 𝜎_𝐸_^2^) is the vector of epistatic genomic marker effects, and 𝜀∼𝑁(0, 𝐼, 𝜎_𝜀_^2^) is the vector of random residual effects. The variance of the additive genomic marker effects (𝜎^2^), epistatic genomic marker effects (𝜎_𝐸_^2^) and random residual effects (𝜎_𝜀_^2^) were summed for the total variance. Each variance component was then expressed as a proportion of the total variance, yielding values between 0 and 1. For the TeoNAM dataset, variance components were calculated per environment and averaged within each population. For the MaizeNAM dataset, variance components were directly calculated at the population level for average performance across environments, since data for each RIL population were not available for each environment. The extended-GBLUP was implemented by ASReml-R (v4.2).

### 4. Genomic prediction model implementation and genomic level analysis

Six individual genomic prediction models were implemented and their inferred marker effects for prediction were extracted using the computational pipeline tool, EasiGP (Tomura et al. 2025a). Following Tomura et al (2025b), three conventional genomic prediction models (ridge regression best linear unbiased prediction (rrBLUP; Meuwissen et al. 2001), BayesB (Meuwissen et al. 2001) and reproducing kernel Hilbert space regression (RKHS; Gianola and Van Kaam 2008)) and three machine learning models (random forest (RF; Breiman 2001), support vector regression (SVR; Drucker et al. 1996) and multi-layer perceptron (MLP; Rosenblatt 1958)) were applied in this study. For rrBLUP, the genomic marker effects are assumed to be normally distributed for all genomic markers. In contrast, some genomic marker effects can be shrunk to zero rather than forming a normal distribution in BayesB (Endelman 2011; de los Campos 2014). BayesB has been widely applied to genomic prediction for crop breeding and hence used as a benchmark model among the Bayesian alphabet models in this study. For RKHS, genomic markers are mapped to the Hilbert space using a kernel that estimates the mean distance between genotypes as a squared Euclidean distance (del los Campos et al. 2009). RF consists of numerous decision trees developed from a fraction of the given dataset. Each decision tree returns prediction values by traversing the tree with input attribute values. The mean predicted value from all the decision trees was calculated as the final prediction value. SVR predicts values by optimising the position of a hyperplane that maximises the number of samples included in the space defined as an epsilon tube. The size of the epsilon tube is determined by the space between the hyperplane and the location of support vectors that create the outer boundary of the tube as the decision boundary (Drucker et al. 1996). Hence, the hyperplane is optimised in consideration of adjusting the position of support vectors. MLP mimics human brain systems by continuous nonlinear aggregation of input data from neighbouring neurons in a forward direction (Dave and Kamlesh, 2014). Those six prediction models have been widely used for genomic prediction in crop breeding and the three prediction models from each category, conventional (including parametric and semi-parametric) and machine learning, were chosen for balanced comparison. The ensemble-average model calculates the mean predicted phenotypes from the predictions returned by the six individual models with equal weights. The genomic prediction models constructed in this study were trained using genomic markers in a numerical format to output predicted phenotypes for each prediction scenario. The details of the prediction scenarios are discussed in “2.6 Model evaluation”.

The following hyperparameter values were assigned to each individual genomic prediction model. For rrBLUP, BayesB and RKHS, the number of iterations was set as 12,000 and the burn-in was 2,000 for the three models in both datasets. Default values were used for all other parameters. For RF and SVR, the default hyperparameter values were employed for both prediction models in both datasets, except for the number of trees (1,000) in RF. MLP was structured with one hidden layer in both datasets. For the TeoNAM dataset, 50 neurons were included in the hidden layer with a dropout of 0 and activated by the Rectified Linear Unit (ReLU; Nair et al. 2010) function. The model was trained for 200 epochs with a learning rate of 0.005 using the Adaptive Moment Estimation with Weight Decay (AdamW; Loshchilov and Hutter 2017) optimiser. For the MaizeNAM dataset, the number of neurons was reduced to 10 with a dropout of 0.1 and also activated by the ReLU function. The model was trained for 2,500 epochs with a learning rate of 0.005 using Root Mean Square Propagation (RMSprop; Tieleman 2012). The selected hyperparameter values reached the highest prediction performance among the combinations in each dataset from our heuristic investigation of optimum hyperparameter value combinations. The detailed associations between the selected hyperparameter values of the individual genomic prediction models and the prediction performance of ensembles can be further investigated as a future research area.

The effect of each marker on trait phenotype prediction was estimated after the development of genomic prediction models as a part of the pipeline in EasiGP. For rrBLUP and BayesB, the allele substitution effects were inferred as marker effects. For RF, marker effects were inferred using feature importance based on the impurity-based approach (Ishwaran 2015). The importance of a feature is determined by the total impurity value derived from the impurity value of each decision node for the feature. Higher importance is allocated to a feature that can clearly divide the data points (RILs) into sub-decision nodes, clustering data points with similar labels (observed phenotypes). For RKHS, SVR, and MLP, Shapley scores (Shapley 1953; Lundberg and Lee 2017) were used as inferred marker effects. Shapley scores in this study estimate the conditional effect of a particular genomic marker. Since Shapley scores are calculated cumulatively by considering possible genomic marker combinations, Shapley scores were also applied to RF at the pairwise level to infer genomic marker-by-marker interaction effects. For the ensemble-average model, the mean values of the normalised marker effects from all the individual genomic prediction models were calculated with the same weights.

The inferred marker effects from each genomic prediction model and the ensemble were mapped to corresponding marker regions using circos plots in the final phase of EasiGP. The generated circos plots also visualised the genome positions of known genes and regulators of DTA and QTL reported from three previous studies. The QTL regions reported by Chen et al. (2019) for the TeoNAM dataset and by Buckler et al. (2009) for the MaizeNAM dataset. Wisser et al. (2019) identified several key genes affecting flowering-time in maize from an empirical selection experiment investigating the short-term evolution of tropical region maize lines. Lastly, Dong et al. (2012) created a list of key gene regions and their interactions for flowering-time in maize. They classified key gene regions into two categories (leaf and shoot apical meristem (SAM)) based on the tissue or organ of the maize plant where they act to determine flowering-time. These genes were also classified into seven categories based on their a priori defined functions (light transduction, clock, photoperiod, autonomous, aging, gibberellin and integrator).

### 5. The Diversity Prediction Theorem

The Diversity Prediction Theorem compares the error of the ensemble-average model against the mean error of individual prediction models in relation to the diversity of the predicted values (Hong and Page 2004; Page 2007; Page 2018):

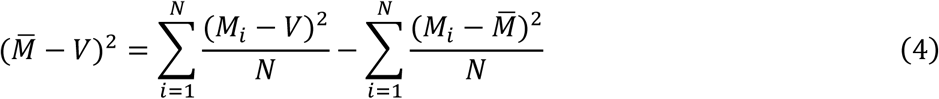

where 𝑀_𝑖_ is the predicted value from the prediction model 𝑖, *M̄* is the mean predicted values from the 𝑖 individual prediction models, 𝑉 is the true value and 𝑁 is the total number of prediction models considered. The Many-Model error (the first term) is calculated by subtracting the prediction diversity (third term) from the average error (second term). In our study, an observed phenotype was defined as the true value 𝑉 and the Many-Model error was regarded as the ensemble error. *M̄* was calculated by averaging the predicted phenotypes from the six individual genomic prediction models used in this study.

Since the scale of phenotypes varied across the traits and datasets, each term value in the Diversity Prediction Theorem was not directly comparable. Hence, the coefficient of variation (CV) was estimated in each term to be used as a metric to measure the diversity level of predicted phenotypes for the ensemble in each prediction scenario:

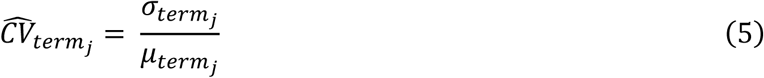

where 𝐶̂𝑉̂_𝑡𝑒𝑟𝑚𝑗_ is an estimate of CV for term 𝑗 in the Diversity Prediction Theorem, the 𝜎_𝑡𝑒𝑟𝑚_𝑗__ represents the standard deviation of the term 𝑗in the theorem and 𝜇_𝑡𝑒𝑟𝑚_𝑗__ represents the mean value of the term 𝑗 in the theorem. A larger estimated CV for the third term of Equation (4) indicates higher diversity in the predicted phenotypes and hence individual genomic prediction models included are considered to be more diverse. The CV was estimated per trait for each dataset.

### 6. Model evaluation

The individual and ensemble genomic prediction models were evaluated by iterative prediction scenarios with different settings. For the TeoNAM dataset, three different training-test set ratios (0.8-0.2, 0.65-0.35 and 0.5-0.5) were applied to mitigate the data size effect. In each population-training-test-set scenario, RIL data were randomly sampled 500 times. Hence, 7,500 prediction scenarios (5 populations * 3 ratios * 500 samples) were developed per trait. The same three training-test set ratios were used in the MaizeNAM dataset. For each of the 25 populations in the dataset, the RILs were sampled 50 times for each population-training-test-set scenario, resulting in 3,750 prediction scenarios (25 populations * 3 ratios * 50 samples) per trait. In each prediction scenario, the genomic prediction models were trained, and their prediction performance was evaluated using the Pearson correlation coefficient and mean squared error (MSE). After recording the prediction performance metrics, the developed genomic prediction models were discarded, treating each prediction scenario as an independent prediction task. Using the recorded predicted phenotypes from the genomic prediction models, the three components of the Diversity Prediction Theorem (Equation (4)) were calculated for the ensemble. At the end of the iterative predictions process, the median values were calculated for both the Pearson correlation coefficients and MSE to develop violin plots, whereas the mean and CV values were calculated for the three terms of the Diversity Prediction Theorem.

## Results

### 1. Two maize NAM datasets and traits showed a distinctive composition of genetic variance components

The estimated genetic variance components for the traits provide a preliminary characterisation of the two target datasets (Fig. 2). For ASI, the median genetic variance component (additive and epistasis) value for the TeoNAM dataset (0.535) was higher than for the MaizeNAM dataset (0.490), indicating that a larger proportion of the phenotypic variance for ASI was explained by genetic effects in the TeoNAM dataset. The median value was used since some populations showed extreme variance component values. This observation is aligned with the genetic diversity previously estimated in the parent lines in the TeoNAM and MaizeNAM datasets; nucleotide diversity (π) across non-overlapping 10-kb windows was higher for the Teosinte lines used in the TeoNAM populations (π = 0.0059; *Z. mays ssp. parviglumis*), compared to the improved maize lines included as parental lines in several MaizeNAM populations (π = 0.0048) (Hufford et al. 2012). In contrast, there was a higher effect of the median genetic variance component for DTA in the MaizeNAM dataset (0.659) compared to the TeoNAM dataset (0.594), indicating that genetic effects explained a larger proportion of the phenotypic variance in the MaizeNAM dataset for DTA. For both datasets, the sum of both additive and epistasis variance components for ASI was smaller than for DTA. This observation likely corresponds to the high environmental plasticity of ASI captured in the residual component (Buckler et al. 2009; Silva et al. 2022). Such a difference in the variance components indicates the potential for distinctive predictive behaviours of the genomic prediction models for the prediction scenarios characterised by traits and datasets, investigated in the following subsections.

**Fig. 2:**
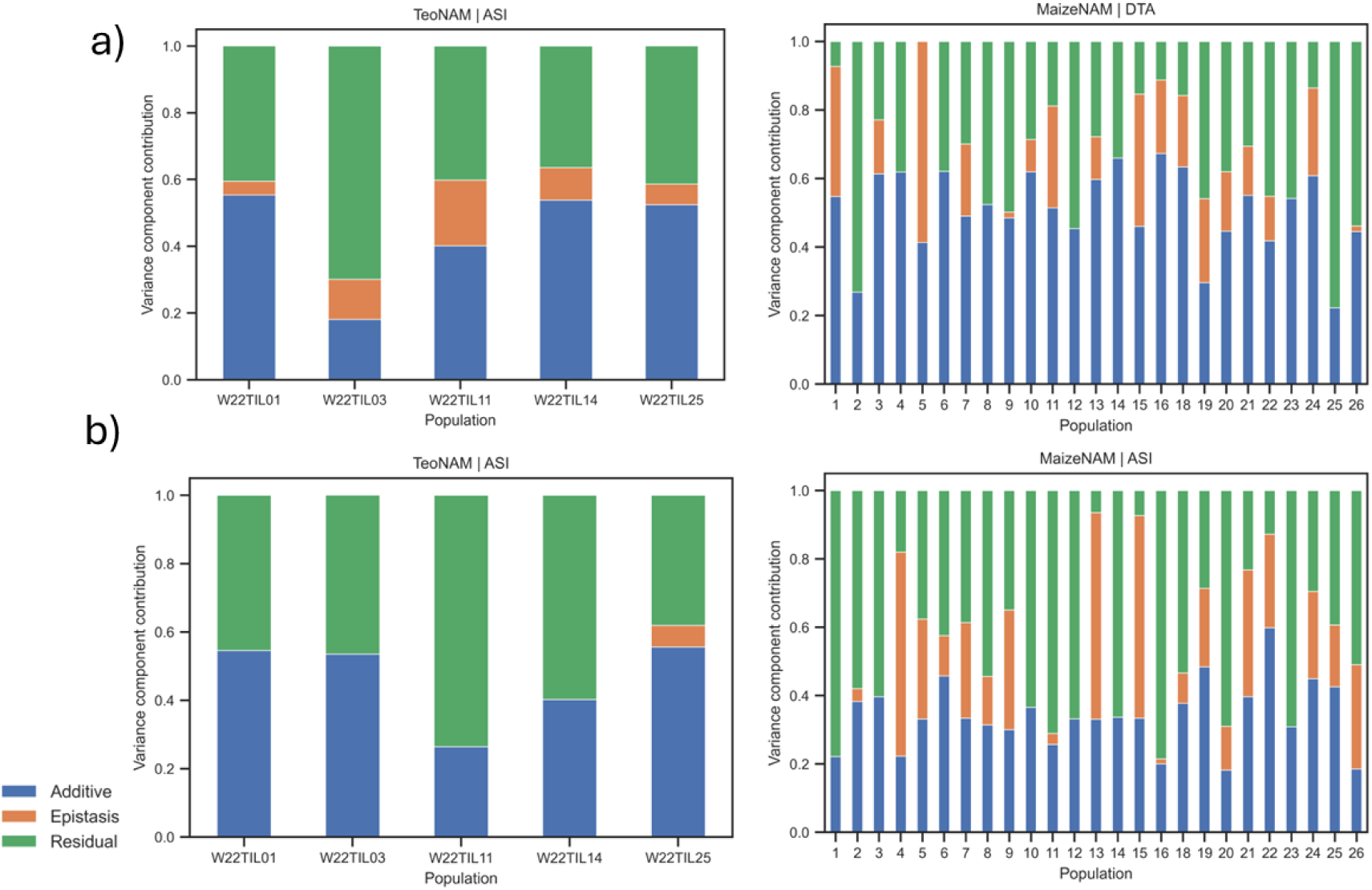
The variance components of each population in the TeoNAM and MaizeNAM datasets for the a) days to anthesis (DTA) and b) anthesis to silking interval (ASI) trait. The relative magnitudes of variance components partitioned into additive (blue), epistasis (orange) and residual (green) effects.

### 2. Ensemble prediction showed improved prediction performance in comparison to the individual genomic prediction models

The ensemble-average model improved the prediction performance across the target traits and datasets relative to the mean prediction performance of individual genomic prediction models by increasing accuracy and reducing error (Fig. 3). For the TeoNAM dataset, the ensemble-average model reached higher median prediction accuracy for DTA (0.842) and ASI (0.505) in contrast to the mean prediction accuracy of the individual genomic prediction models for DTA (0.741) and ASI (0.473). The ensemble-average model also had a smaller median prediction error for DTA (the ensemble-average model = 11.033 and the mean of individual genomic prediction models = 14.699) and ASI (the ensemble-average model = 4.339 and the mean of individual genomic prediction models = 4.638). For the MaizeNAM dataset, the median prediction accuracy for DTA and ASI was higher for the ensemble-average model (0.640 and 0.464, respectively) compared to the mean performance of the individual genomic prediction models (0.598 and 0.432, respectively). Similarly, there was improvement in the median prediction error for both traits (DTA = 3.471 and ASI = 0.980 in contrast to DTA = 4.467 and ASI = 1.023). This consistent outperformance of the ensemble-average model was also observed in the predictions using BLUPs for the MaizeNAM dataset, showing a similar median prediction performance for the BLUEs in both traits (Fig. S3). The observed results indicated that the ensemble-average model consistently outperformed the mean prediction performance of the individual models.

**Fig. 3:**
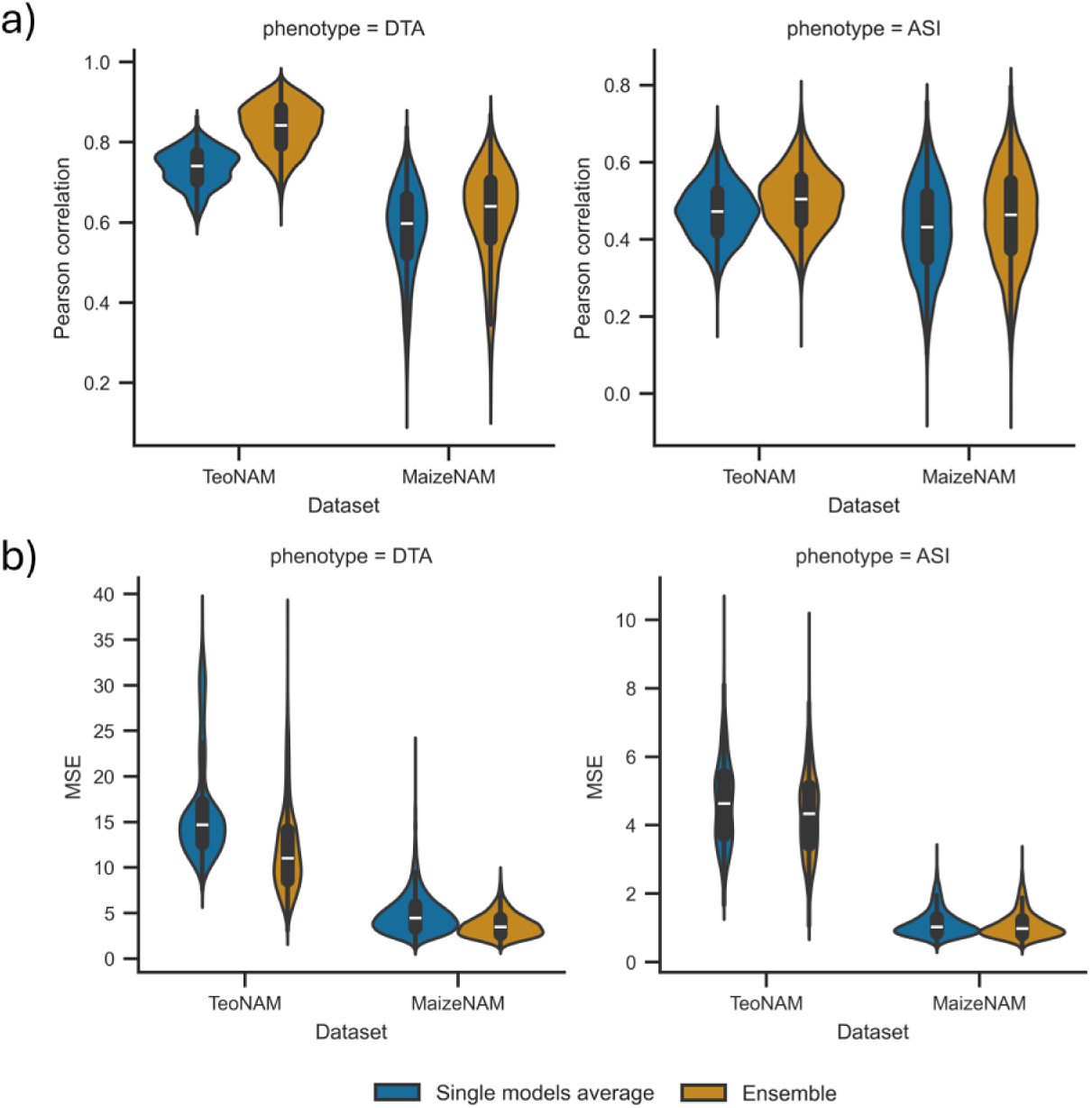
The comparison of prediction performance between the average performance of the individual genomic prediction models (Single models average) and the ensemble-average model (Ensemble) for the days to anthesis (DTA) and anthesis to silking interval (ASI) traits in the TeoNAM and MaizeNAM datasets. The prediction performance was measured using the two metrics: a) Pearson correlation and b) mean squared error (MSE). The prediction performance was measured in 7,500 prediction scenarios for the TeoNAM dataset and 3,750 prediction scenarios for the MaizeNAM dataset for each trait. The prediction scenarios were generated by the combination of the three training-test ratios (0.8-0.2, 0.65-0.35 and 0.5-0.5), populations and sampling numbers. The width of the violins represents the distribution of performance metrics. The white horizontal lines on the black box plots show the median value for each metric. The whiskers extend 1.5 times the interquartile range.

Higher prediction performance of the ensemble-average model was also observed in the comparison with the individual genomic prediction models (Fig. S4), achieving the highest prediction performance or equivalent performance to the best individual genomic prediction models throughout the trait-by-dataset prediction scenarios in this study. The ensemble-average model was selected as one of the top three best genomic prediction models, competing with rrBLUP, BayesB and RKHS, in the range of generated prediction scenarios from this study. When the prediction performance of the genomic prediction models was analysed at the population level in each dataset (Table S2), the ensemble also showed stable improved prediction performance by being included as one of the top three prediction models in trait-by-population prediction scenarios. For prediction accuracy, the ensemble showed the highest percentage (78.3%) of achieving the top three most accurate prediction models measured in terms of Pearson correlation, followed by BayesB (75%) and rrBLUP (73.3%). For the prediction error, rrBLUP and BayesB reached the highest percentage of achieving the top three lowest median MSE (78.3%), followed by RKHS (68.3%) and ensemble (65.0%). In contrast, the machine learning models (RF, SVR and MLP) remained at lower prediction performance throughout the trait-by-population prediction scenarios in both prediction accuracy and error by consistently having a low percentage (less than 10.0%). The stability in the improved prediction performance of the ensemble was also observed in trait-by-training set ratio prediction scenarios in each dataset (Fig. S5), showing constant performance reduction rate in proportion to the reduction in training set size along with rrBLUP, BayesB, RF and SVR. The ensemble minimised the effect of the rapid performance reduction that was observed for RKHS and MLP, attributed to the training set size reduction in some prediction scenarios. The comparison with the individual genomic prediction models demonstrated that the ensemble-average model consistently reached a higher prediction accuracy across a wide range of prediction scenarios.

### 3. Diversity contributed to the performance improvement of the ensemble

The level of prediction improvement of the ensemble varied with traits and datasets, associated with the level of diversity in predicted phenotypes. The CV for the third term was higher in the TeoNAM dataset for both traits (DTA = 1.99 and ASI = 2.15) in contrast to the MaizeNAM dataset (DTA = 1.80 and ASI = 1.15) (Table 2). A higher CV for the third term indicates that the predicted phenotypes from the individual genomic prediction models are more diverse. Since greater improvements in prediction performance were observed in the TeoNAM dataset compared to the MaizeNAM dataset, the higher CV values for both DTA and ASI in the TeoNAM dataset were associated with higher diversity in the predicted phenotypes from the individual genomic prediction models. This finding indicated that there was a positive association between the level of diversity in the predicted phenotypes and the prediction performance of the ensemble.

**Table 2:**
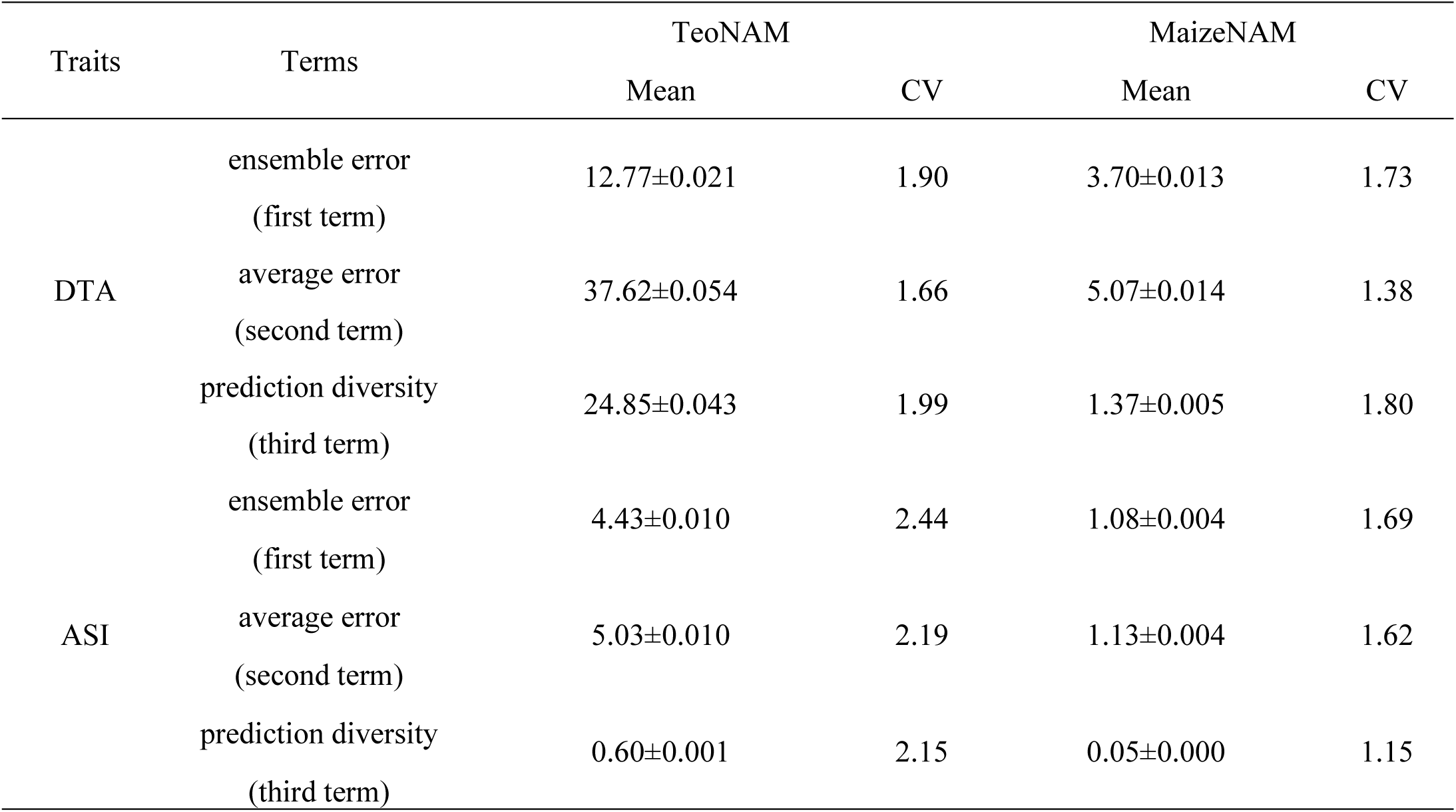
Measurement of diversity in individual genomic prediction models included in the ensemble models using the framework of the Diversity Prediction Theorem. The mean value of each term in the Diversity Prediction Theorem was calculated for the days to anthesis (DTA) and anthesis to silking interval (ASI) traits in the TeoNAM and MaizeNAM datasets. The ensemble error, average error and prediction diversity represent the first, second and third terms in the theorem, respectively. In each dataset, “Mean” represents the actual mean term value, whereas “CV” indicates the coefficient of variation for each term value. The sign “±” indicates the standard error of each corresponding mean value.

The comparison of the predicted DTA at both the phenotype and genome marker levels indicated a different degree of information diversity between the datasets and between prediction model groups, conventional (rrBLUP, BayesB and RKHS) versus machine learning (RF, SVR and MLP) (Fig. 4). Pairwise comparisons between the conventional models and machine learning models in the MaizeNAM dataset demonstrated stronger positive associations (mean predicted phenotypes: *r* = 0.961 and mean inferred marker effects: *r* = 0.431) compared to the TeoNAM dataset (mean predicted phenotypes: *r* = 0.165 and mean inferred marker effects: *r* = 0.603). These results indicate that, for the TeoNAM dataset, the individual genomic prediction models returned more diverse predicted phenotypes, consistent with the substantial variation in marker effect estimates between the prediction model groups. The relatively high diversity in the TeoNAM dataset was also observed for ASI at the phenotype levels (Fig. S6). The MaizeNAM showed a higher positive association at the phenotypic level (mean predicted phenotypes: *r* = 0.954 and mean inferred marker effects: *r* = 0.461) compared to the TeoNAM dataset (mean predicted phenotypes: *r* = 0.197 and mean inferred marker effects: *r* = 0.750). This general trend of strong positive associations at the phenotype and genome levels in both traits for the MaizeNAM dataset was also illustrated in the pairwise comparison results for the six individual prediction models in both datasets (Fig. S7, S8; Table S3, S4). The only exception was observed in the pairwise comparison with RKHS at the genome level for the MaizeNAM dataset, which warrants further investigation into the underlying cause in future studies. This higher diversity among the prediction models at both phenotype and genome levels for the TeoNAM dataset likely contributed to the higher level of improvement in prediction performance of the ensemble model in comparison to the MaizeNAM dataset (Fig. 3, 4).

**Fig. 4:**
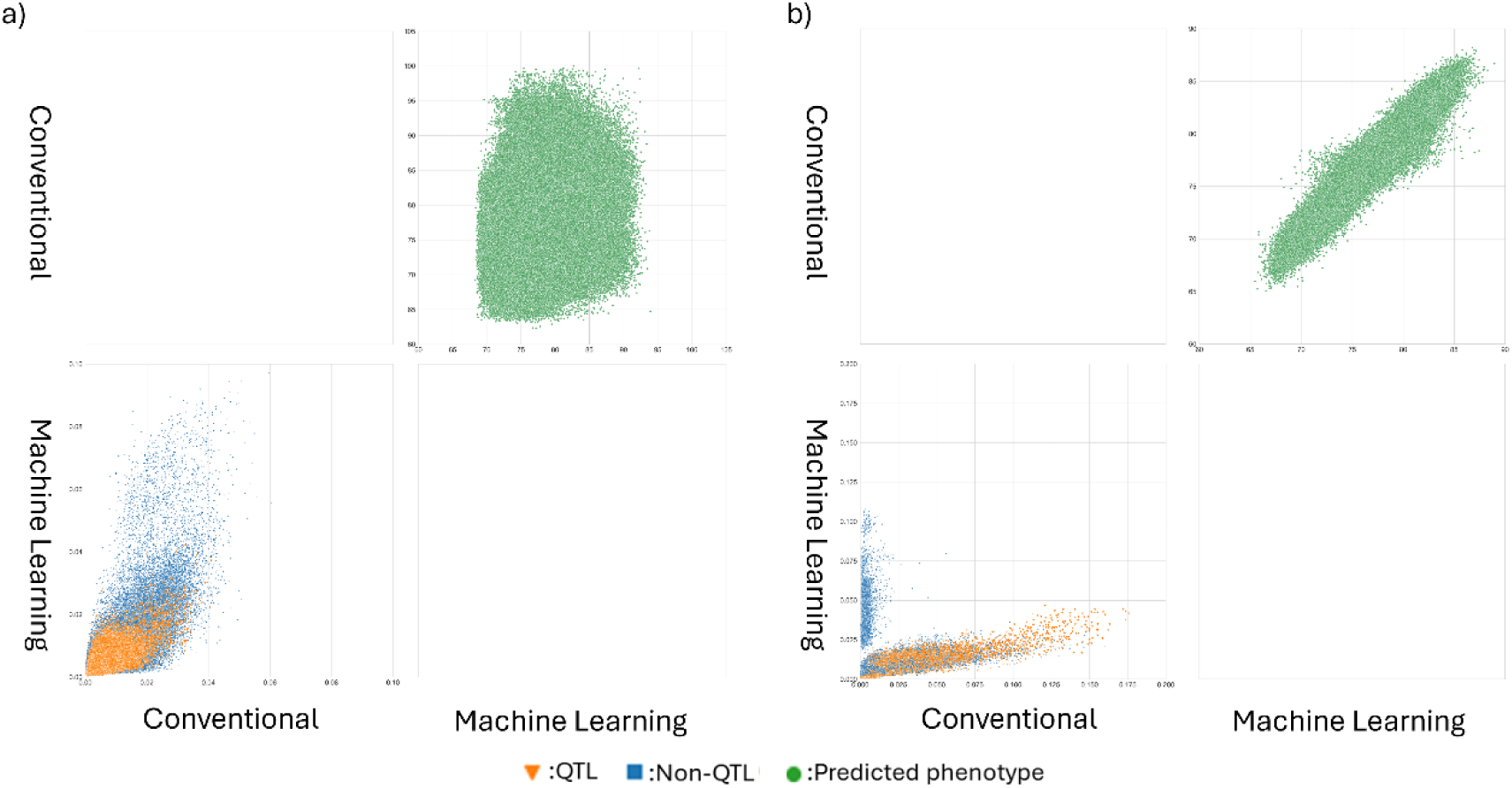
Pairwise comparisons of the conventional (rrBLUP, BayesB and RKHS) and machine learning models (RF, SVR and MLP) for the days to anthesis (DTA) trait across all the prediction scenarios for the a) TeoNAM (7,500 prediction scenarios) and b) MaizeNAM (3,750 prediction scenarios) datasets. The prediction scenarios were generated by the combination of the three training-test ratios (0.8-0.2, 0.65-0.35 and 0.5-0.5), populations and sampling numbers. The genomic prediction model groups were compared for mean predicted phenotypes (top right triangle) and mean normalised genomic marker effects (the bottom left triangle) calculated within each prediction model category. The green dots represent a pair of predicted phenotypes of RIL samples in the test sets for each prediction scenario. The blue squares and orange triangles represent a pair of extracted genomic marker effects from each genomic marker in each sample scenario that were identified as non-QTL and QTL markers, respectively, by Chen et al. (2019) for the TeoNAM dataset and Buckler et al. (2009) for the MaizeNAM dataset.

There were some important differences observed between the conventional and machine learning prediction models at the level of the estimated marker effects. The effects for a significant fraction of the markers were reduced to zero for the conventional prediction models due to the shrinkage factors incorporated in their prediction mechanisms, while the assigned effects for these markers from the machine learning models did not show such a shrinkage. Consequently, the genomic marker effects were not highly correlated between the conventional and machine learning model groups. This trend was especially emphasised for the MaizeNAM dataset. The comparison of the marker effects between the two prediction model groups clearly indicated that many genomic marker effects were captured differently as features between the two prediction model groups based on their unique algorithms.

The diversity of the prediction models at the genome level was further investigated by mapping estimated marker effects from each to corresponding genome regions for both traits (Fig. 5, S9). The highlighted genomic markers and predictive patterns from each genomic prediction model were more variable in the TeoNAM dataset (Fig. 5a, S9a) in comparison to the MaizeNAM dataset (Fig. 5a, S9b), consistent with the expectations of the Diversity Prediction Theorem framework (Table 2) and the results of the ensemble model prediction (Fig. 3). While the different genomic prediction models identified many similar regions of the genome, the effect of each marker identified in these regions varied considerably in the TeoNAM dataset (Fig. 5a, S9a). In the MaizeNAM dataset, on the other hand, there was more consistency across the genomic prediction models (Fig. 5b, S9b). The genomic prediction models detected similar genomic regions with a stronger consensus on the effect of each marker. This observation also highlights the positive association between the prediction performance of the ensembles and the diversity of individual prediction models included in the ensembles at the genome level.

**Fig. 5:**
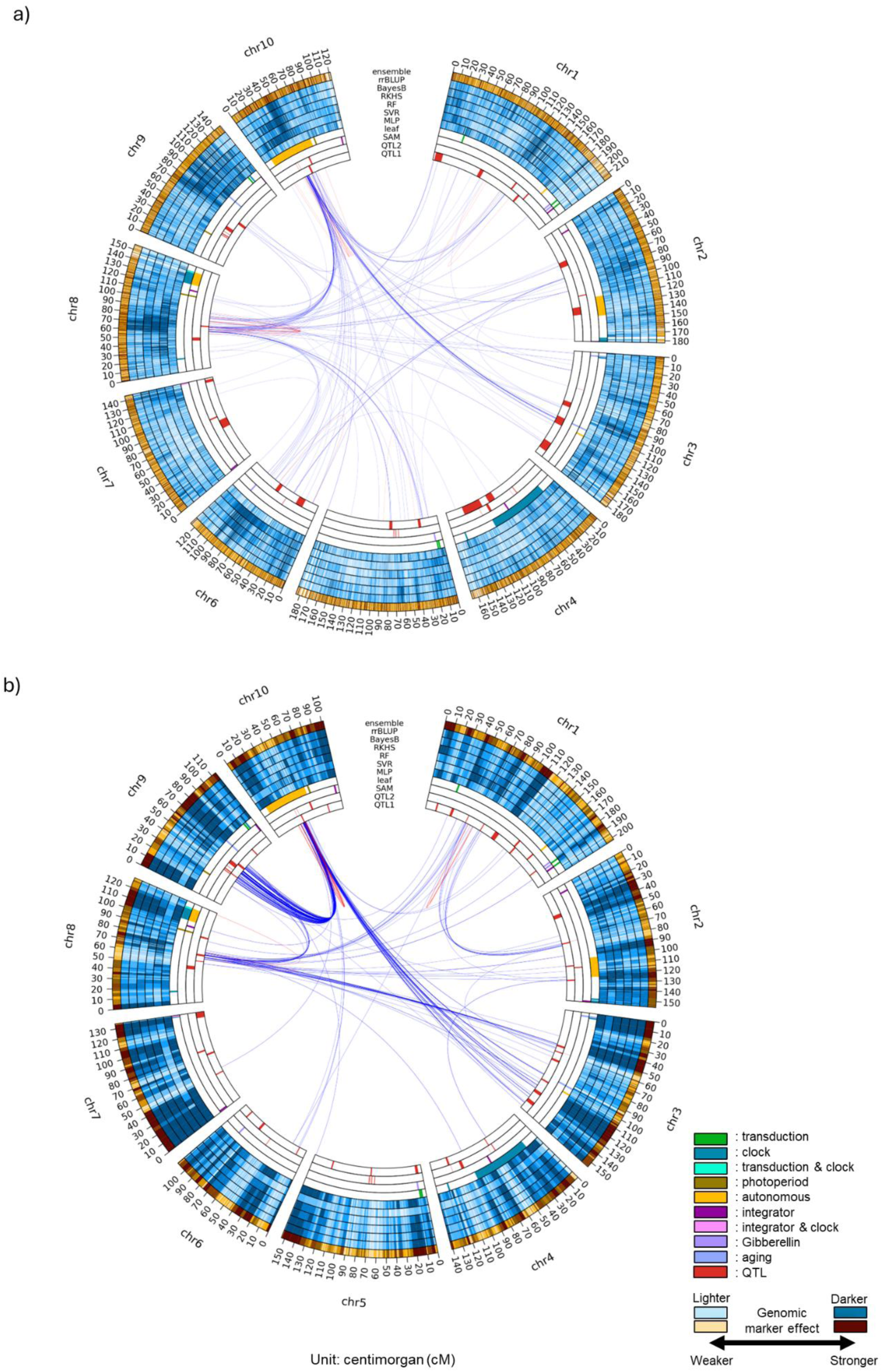
Circos plots for the days to anthesis (DTA) trait using the a) TeoNAM and b) MaizeNAM datasets. The innermost (QTL 1) ring shows the QTL gene regions estimated by Chen et al. (2019) for the TeoNAM dataset and by Buckler et al. (2009) for the MaizeNAM dataset. The second innermost ring (QTL 2) represents the QTL gene regions identified by Wisser et al. (2019). The third and fourth innermost rings represent gene regulators that affect the shoot apical meristem (SAM) and leaf, respectively, identified by Dong et al. (2012). The blue rings (fifth to tenth) represent the genomic marker effects across the gene regions estimated by MLP, SVR, RF, RKHS, BayesB and rrBLUP, respectively. The outermost orange ring is the genomic marker effects for the ensemble-average model (ensemble). The numbers at the outermost ring represent genetic distance in centimorgans (cM). The darkness level of the blue and orange colours indicates the strength of the genomic marker effects, sectioned into ten levels using the quantiles. Darker colours represent higher genomic marker effect levels. The red and blue lines between genome regions are the genomic marker interaction effects calculated by pairwise Shapley scores from RF (top 0.01%; red = within chromosome and blue = between chromosomes).

It is also noted that the individual genomic prediction models combine genomic marker effects from both QTL and non-QTL regions to predict target trait phenotypes rather than only using the estimated marker effects from QTL regions identified in the previous analyses of both datasets (Buckler et al. 2009; Chen et al. 2019). For the TeoNAM dataset, while the markers at the peaks of the QTL identified by Chen et al. (2019) were included in the genomic prediction models, strong weights were less frequently assigned by the genomic prediction models (Fig. 4a, S6a, S7a, S8a; lower triangle of pairwise comparisons). For the MaizeNAM dataset, in contrast, while the genomic prediction models allocated stronger weights to the markers at the peaks of the QTL identified by Buckler et al. (2009) from their analysis of the same dataset, strong marker effects were also assigned to several markers corresponding to non-QTL regions (Fig. 4b, S6b, S7b, S8b; lower triangle of pairwise comparisons). The individual genomic prediction models not only utilised the association between QTL regions and corresponding traits to predict phenotypes but also included non-QTL regions as features by assigning stronger effect weights to them.

### 4. Several genome regions containing key gene regulators were highlighted by EasiGP

The constructed circos plots showed several genome regions with overlap between the genomic marker regions highlighted by the genomic prediction models and known key gene regulators for the traits (Fig. 5). For DTA, the features of the genomic prediction models were associated with genome regions containing several key maize flowering-time genes in both datasets. For example, in the TeoNAM dataset, the region between 35 cM and 50 cM in chromosome 10 was repeatedly targeted across genomic prediction models (with MLP as an exception). This region contains *ZmCCT10*, involved in regulating flowering-time in maize, known to upregulate the circadian clock as part of the photoperiod pathway (Dong et al. 2012; Chen et al. 2019; Wisser et al. 2019). The region containing *ZmCCT10* also showed strong interactions with the region in chromosome 8 (between 60 cM and 80 cM). This region includes *ZCN8*, a well-established gene controlling flowering-time in maize that is known to interact with *ZmCCT10* (Dong et al. 2012; Chen et al. 2019; Wisser et al. 2019). In the MaizeNAM dataset, in addition to the same two regions identified in the TeoNAM dataset, the genomic prediction models identified another region between 50 cM and 60 cM in chromosome 9 that strongly interacted with the region containing *ZmCCT10* in chromosome 10 and the region containing *ZCN8* in chromosome 8. This chromosome 9 region contains *ZmCCT9* (Wisser et al. 2019), another gene involved in the photoperiod pathway that negatively regulates *ZCN8* (Huang et al. 2018). For ASI, several examples of overlap between the regions highlighted by the genomic prediction models and previously identified QTL regions (Buckler et al. 2009; Chen et al. 2019) were also observed in both datasets (Fig. S9). Such examples of overlap between the regions of the genome identified across genomic prediction models and the key gene regulators from the literature in both datasets indicated that these genomic prediction models repeatedly identified regions containing several known causative loci. The stronger consensus in the highlighted feature of marker effects was also observed in the results based on the analyses conducted on BLUPs in the MaizeNAM dataset, with minor differences in the extracted genomic marker-by-marker interactions (Fig. S10, S11). Combined with the observed results from the previous subsection, genomic-level analysis of the prediction models revealed that they repeatedly identified several regions of the genome containing key genes for maize flowering-time traits, while the diverse effects of markers across other genome regions contributed to the predictive performance of the ensembles.

## Discussion

Breeders have continuously searched for prediction methods that can consistently and accurately distinguish individuals with the desired traits in their breeding programs (Cooper et al. 2014a,b; Voss-Fels et al. 2019; Diepenbrock et al. 2022; Crossa et al. 2025; Escamilla et al. 2025). The results from previous genomic prediction research indicate that genomic prediction models can accurately identify superior individuals under certain prediction scenarios (Azodi et al. 2019; Diepenbrock et al. 2022). However, a priori, it is often not possible to obtain information that determines the optimal combination of scenarios and genomic prediction models without a significant trial-and-error process, since no trends or patterns in the combinations are often observed (Jeon et al. 2023; Crossa et al. 2025). For instance, RKHS reached the highest prediction accuracy on wheat yield prediction for one wheat dataset (Heslot et al. 2012). However, this trend may not always be observed. In our study, rrBLUP and BayesB showed relatively high prediction performance across different prediction scenarios. The prediction performance superiority for RKHS was not consistent, showing diminished RKHS prediction accuracy at one of the lowest levels for wheat yield traits in another dataset (Heslot et al. 2012). This trend of inconsistency in the best genomic prediction model was observed in other studies as well (Endelman 2011; Meher et al. 2022; Lourenço et al. 2024; Montesinos-López et al. 2024). The overwhelming empirical evidence strongly supports the conclusion of no consistently outperforming individual genomic prediction models.

Machine learning models also failed to consistently improve the prediction performance despite their ability to detect nonlinear prediction patterns, as observed in this study. Heslot et al. (2012) also emphasised lower prediction performance of machine learning models, including RF, SVR and MLP, compared to conventional genomic prediction models such as rrBLUP and Bayesian approaches for several yield traits in wheat and maize. Their results aligned with the observed results in our study, showing poor prediction performance of RF, SVR and MLP when used individually. Machine learning approaches did not necessarily outperform the conventional genomic prediction models despite the expectation of outperformance via capturing complex nonlinear prediction patterns incorporated in datasets (Howard and Lipka 2025). A smaller total number of data points (RIL records) in relation to the total number of included genomic markers might have hindered the detection of key prediction patterns, often described as the curse of dimensionality (Bellman 1957). However, MLP reached higher prediction performance than GBLUP in other prediction datasets and traits, including yield traits, in wheat, maize and rice (Montesinos-López et al. 2025). The current machine learning models can outperform the conventional genomic prediction models for some prediction scenarios, drawing the same conclusion that the most preferable genomic prediction model is highly scenario dependent.

In contrast, the ensemble of multiple individual genomic prediction models consistently improved prediction performance in accordance with previous investigations (Kick and Washburn 2023; Washburn et al. 2025) and the expectations of the Diversity Prediction Theorem, given that the individual genomic prediction models included in the ensembles were diverse (Hong and Page 2004; Page 2018; Messina et al. 2025). Hence, the improvement in prediction performance achieved by the ensemble was marginal in some cases, depending on the diversity level in the predictive information included in the individual prediction models. However, the performance of the ensemble approach was always at least comparable to the mean prediction performance of individual genomic prediction models. The significance of our study arises from connecting the theorem to the empirical investigation, showing that the effect of the theorem is applicable in actual genomic prediction settings. The ensemble of predicted phenotypes through the mean value calculation consistently reduced prediction error for other genomic prediction studies in wheat (Wallach et al. 2018), maize (Kick and Washburn 2023; Tomura et al. 2025b) and coffee (Nascimento et al. 2024) or reached the equivalent performance to the best multi-kernel RKHS models in wheat and pig traits (Tusell et al., 2014), indicating that the prediction performance of the mean value-based ensemble models in their studies reached the highest or equivalent to that of the best individual prediction model. Hence, both theory and the body of empirical evidence indicate that an ensemble of genomic prediction models may be a more efficient approach than continuous trial-and-error investigation to determine the best individual genomic prediction model for each trait and dataset.

The success of the ensemble approach demonstrated in this study is interpreted in part to be a consequence of the high levels of genetic diversity represented in both datasets (Buckler et al. 2009; Hufford et al. 2012; Chen et al. 2019). The high levels of genetic variation in both NAM experiments contributed to the high levels of diversity in the predicted phenotypes and the estimated genomic marker effects from the individual genomic prediction models. While high diversity in predicted values can improve the performance of the ensemble approach, excessively diverse prediction values may not always lead to further increases in the prediction performance. The level of diversity needs to be constrained within a certain range in relation to the prediction performance of individual prediction models that vary by prediction scenarios (Bian and Chen 2021; Wood et al. 2023). The benefits of such constrained levels of diversity align with the implications of the theoretical framework of the Diversity Prediction Theorem (Page 2018). This framework can be applied to investigate the influence of diverse predicted values (Equation (4) third term) towards the prediction error of the ensemble (Equation (4) first term) in relation to the mean predicted errors (Equation (4) second term) of individual prediction models, as was conducted for the TeoNAM and MaizeNAM datasets here.

In addition to the contribution to performance improvement through the higher prediction accuracy and lower prediction error, ensemble approaches also enabled the investigation of the repeatability in the highlighted genomic marker effects included as features in the genomic prediction models. Although the six prediction models utilised unique prediction algorithms, ranging from the infinitesimal additive models with shrinkage factors to complex nonlinear prediction mechanisms, the constructed prediction models repeatedly highlighted several genome regions where prior evidence indicates the presence of key genes for control of maize flowering-time. This high repeatability showed that prediction models successfully assigned large genomic marker effects within regions containing known genes, interpreted to contribute to increasing their prediction performance. Through combining the genomic prediction models into an ensemble, it is interpreted that the different models capture a larger spectrum of the potential trait genetic variation for the reference population of genotypes in the TeoNAM and MaizeNAM studies. A more comprehensive representation of the trait genetic variation by the ensembles is interpreted to have also contributed to their improved prediction performance. The genome regions with high repeatability across genomic prediction models and the TeoNAM and MaizeNAM datasets provide targets for further investigation to confirm the influences of known genes and potentially discover new genes involved in determining trait genetic variation. The use of a simulation dataset can be a potential method to support further investigations. For example, by predefining QTL regions, the reliability of genome regions highlighted by the ensembles can be quantitatively analysed by calculating the precision of overlaps between the highlighted and true QTL regions. Reliability can also be investigated in relation to the association with the complexity of the trait genetic architecture embedded in a simulated dataset. Such a comparative analysis can comprehensively assess the reliability of repeatedly highlighted genome regions by ensembles as a future research target.

While the ensemble of diverse individual genomic prediction models can be a promising method for consistent prediction performance improvement with the capture of the trait genetic variation, the operational time to compute the ensemble can be an important factor impacting the practicality of the ensemble for large scale operations in breeding programs. Assuming that input data are preprocessed through imputation and genomic marker filtering, the most time-consuming process can be the development of the input matrix for the ensemble from the predicted phenotype vector of each individual genomic prediction model to apply an efficient mean calculating operation. This process can also be error-prone since the order of data points (RILs) for the predicted phenotypes must be sorted to be identical across all the individual genomic prediction models. This problem can be simplified by the development of a computational pipeline that can automate all tasks involved in the ensemble operation. Using a computational tool such as EasiGP (Tomura et al. 2025a), predicted phenotypes are extracted from user-selected individual genomic prediction models and converted into an input matrix for the ensemble automatically, removing the manual operations once the initial input genomic marker data are provided. The improvement of such ensemble computational tools can be operationalised to help breeders apply the ensemble approach to their large-scale breeding programs within the time requirements for applications.

Given that prediction performance improvement was observed for the simplest ensemble approach (averaging predicted phenotypes with equal weights), which was applied herein to demonstrate the methodology, there is potential for further performance improvement using advanced ensemble design algorithms. One promising approach can be weight optimisation. The minimisation of the Many-Model error in the Diversity Prediction Theorem (Page 2018) can be employed as one method to optimise the weights, since the Many-Model error represents the prediction error of the ensemble. This and other forms of ensemble optimisation are an active area for further research. The integration of prior knowledge into ensembles can be another approach to potentially improve the prediction performance of ensembles. Our investigation of the TeoNAM and MaizeNAM datasets highlighted the effects of several known key flowering-time genes (*ZmCCT9*, *ZmCCT10* and *ZCN8*) and emphasised the importance of other previously unknown genomic regions. Extracted predictive patterns from interpretable genomic prediction models can be utilised as data-driven prior knowledge to supervise the training of genomic prediction models. Prior knowledge can provide prediction models with external information to integrate natural biophysical laws within their prediction mechanisms (Technow et al. 2015; Messina et al. 2018, 2025; Von Rueden et al. 2021; Diepenbrock et al. 2022; Cooper et al. 2025). Such applications of prior knowledge can mitigate the risk of overfitting prediction models, especially when the training set is small and hence the prediction models cannot be well-trained only using the training set (Schapire et al. 2002). The prediction performance of individual genomic prediction models is expected to increase and, consequently, improve the prediction performance of ensembles of these models.

Overall, our experimental investigation was consistent with previous studies showing that an ensemble of multiple genomic prediction models can improve prediction accuracy, indicating its potential to accelerate genetic gain. In addition to demonstrating an effective approach to achieve high prediction performance across multiple dataset scenarios, the ensemble-based methods can be applied to highlight key genomic regions contributing to the standing genetic variation for traits as demonstrated here for the TeoNAM and MaizeNAM datasets. Such improvements in prediction performance motivate further consideration of ensemble-based prediction methodology for breeding applications. Future studies will explore opportunities to extend the ensemble-average model used in this study, including the assessment of prediction performance improvement by optimising weightings of models and integrating prior knowledge (Diepenbrock et al. 2022; Messina et al. 2025).

## Data and code availability

The TeoNAM dataset (Chen et al. 2019) used in this study is located at https://datacommons.cyverse.org/browse/iplant/home/shared/panzea/genotypes/GBS/TeosinteNAM for the genotypes and https://gsajournals.figshare.com/articles/dataset/Supplemental_Material_for_Chen_et_al_2019/9250682 for the phenotypes. The MaizeNAM dataset (Buckler et al. 2009) used in this study is located at https://cbsusrv04.tc.cornell.edu/users/panzea/download.aspx?filegroupid=10 for the genotypes and phenotypes. The computational tool used in this study and the downloaded datasets are located at https://github.com/ShunichiroT/EasiGP.

## Author contribution

ST designed the experiment, analysed data, visualised the results and wrote the manuscript. OP and MJW provided feedback on the analysis, results and manuscript draft. MC formalised the concept, introduced the datasets, designed the experiment and provided feedback on the analysis, results and manuscript draft. All authors discussed the results.

## Acknowledgments

We thank the National Computational Infrastructure (NCI) and the Research Computing Centre (RCC) at the University of Queensland for providing access to the High Performance Computing (HPC) machines. We also thank Dr Seema Yadav for the support in the variance component analysis.

---

**Table S1:** The final number of recombinant inbred line (RIL) records and genomic markers (SNPs) in each population of the TeoNAM and MaizeNAM dataset.

| Dataset | Population | RIL records | SNP numbers |
| --- | --- | --- | --- |
| TeoNAM | W22TIL01 | 444 | 274 - 322 |
|  | W22TIL03 | 540 | 295 - 342 |
|  | W22TIL11 | 438 | 268 - 314 |
|  | W22TIL14 | 460 | 270 - 320 |
|  | W22TIL25 | 616 | 294 - 341 |
| MaizeNAM | 1 | 194 | 170 - 201 |
|  | 2 | 196 | 177 - 206 |
|  | 3 | 191 | 217 - 251 |
|  | 4 | 196 | 188 - 211 |
|  | 5 | 187 | 209 - 235 |
|  | 6 | 185 | 199 - 222 |
|  | 7 | 193 | 183 - 213 |
|  | 8 | 196 | 206 - 236 |
|  | 9 | 189 | 197 - 225 |
|  | 10 | 192 | 192 - 224 |
|  | 11 | 194 | 186 - 215 |
|  | 12 | 193 | 190 - 218 |
|  | 13 | 126 | 199 - 231 |
|  | 14 | 196 | 182 - 213 |
|  | 15 | 185 | 185 - 210 |
|  | 16 | 194 | 175 - 200 |
|  | 18 | 192 | 176 - 199 |
|  | 19 | 196 | 183 - 206 |
|  | 20 | 188 | 197 - 228 |
|  | 21 | 184 | 190 - 214 |
|  | 22 | 193 | 188 - 215 |
|  | 23 | 181 | 174 - 199 |
|  | 24 | 183 | 183 - 210 |
|  | 25 | 188 | 201 - 227 |
|  | 26 | 187 | 202 - 230 |

**Table S2:**
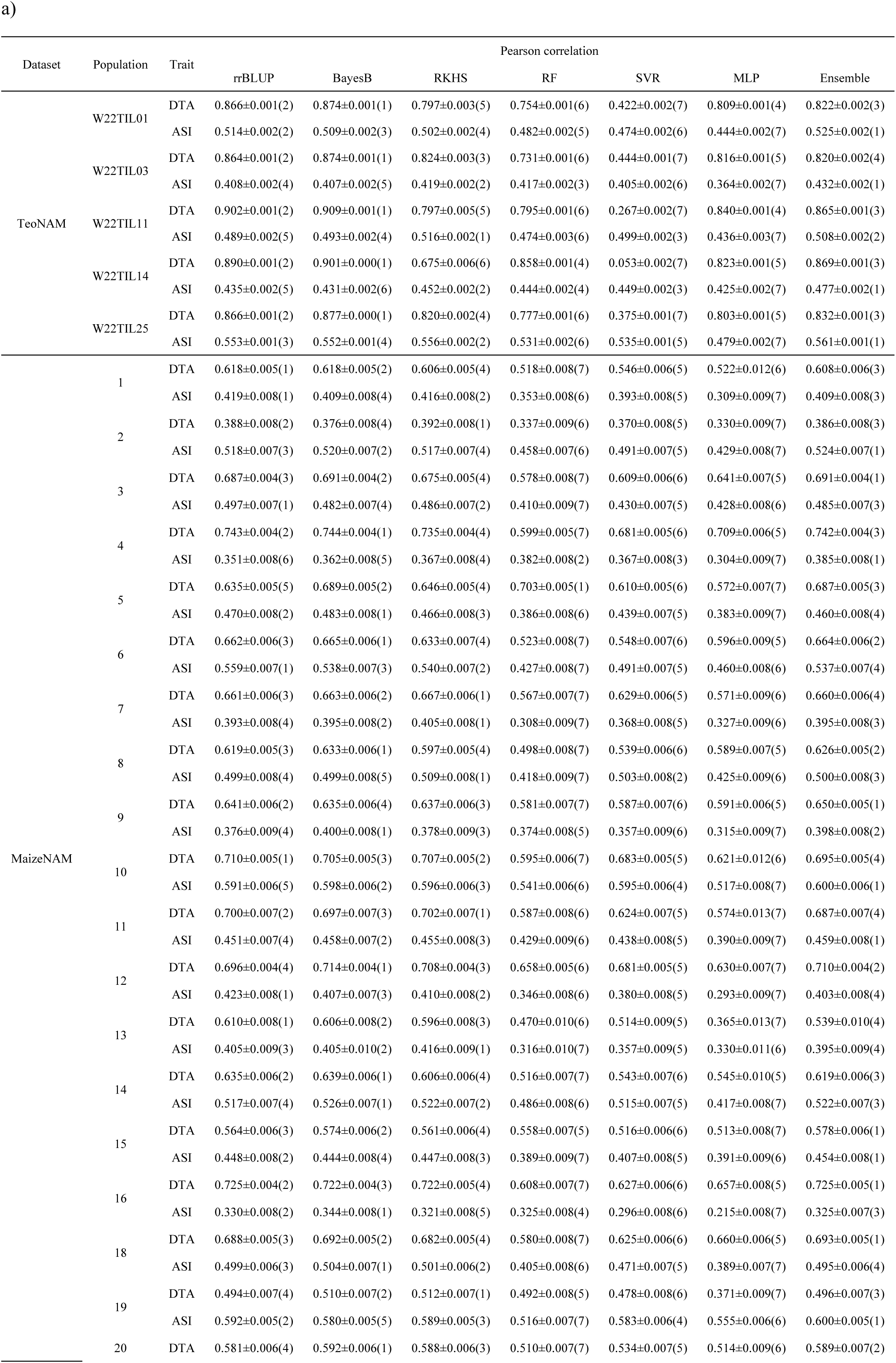
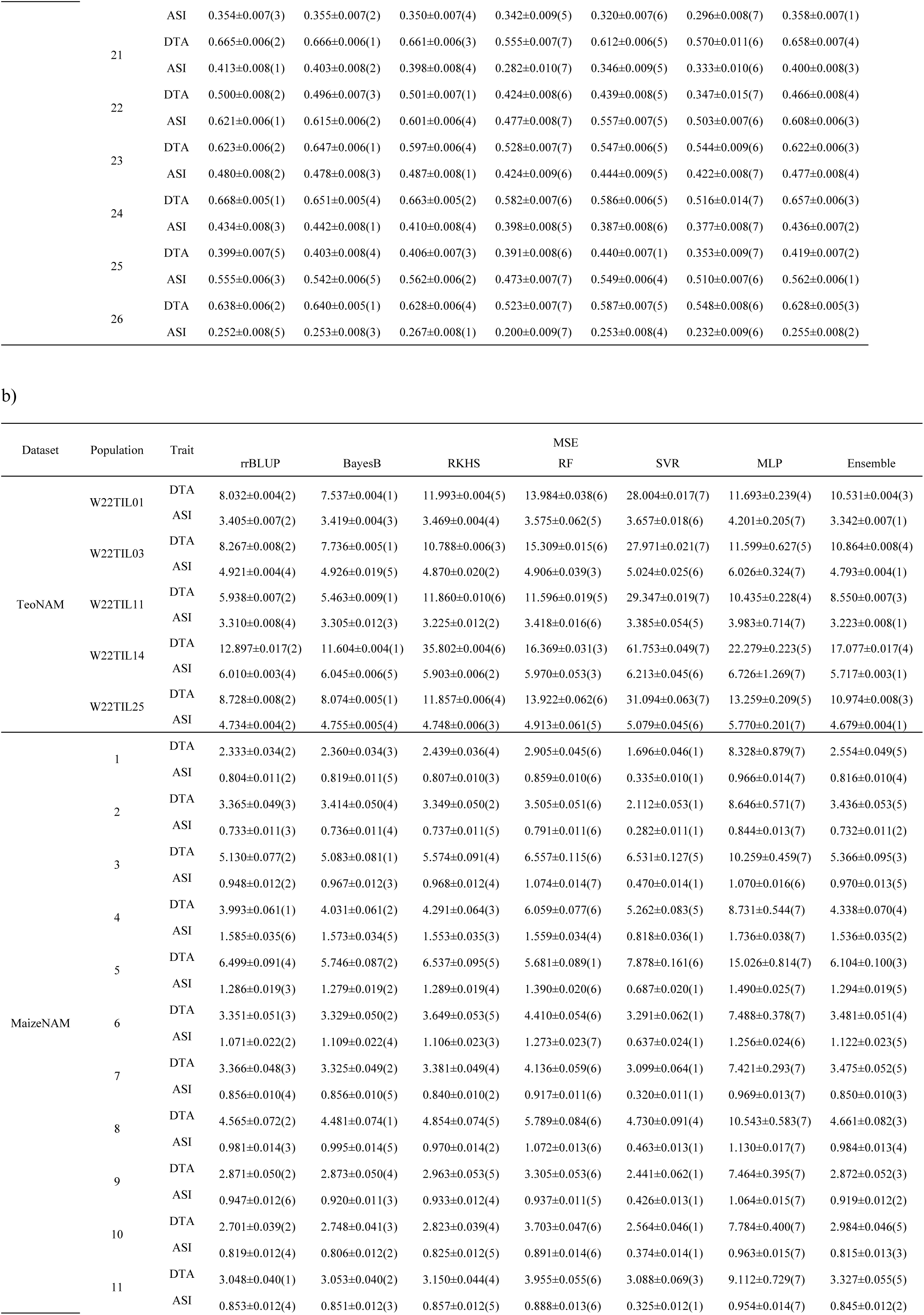
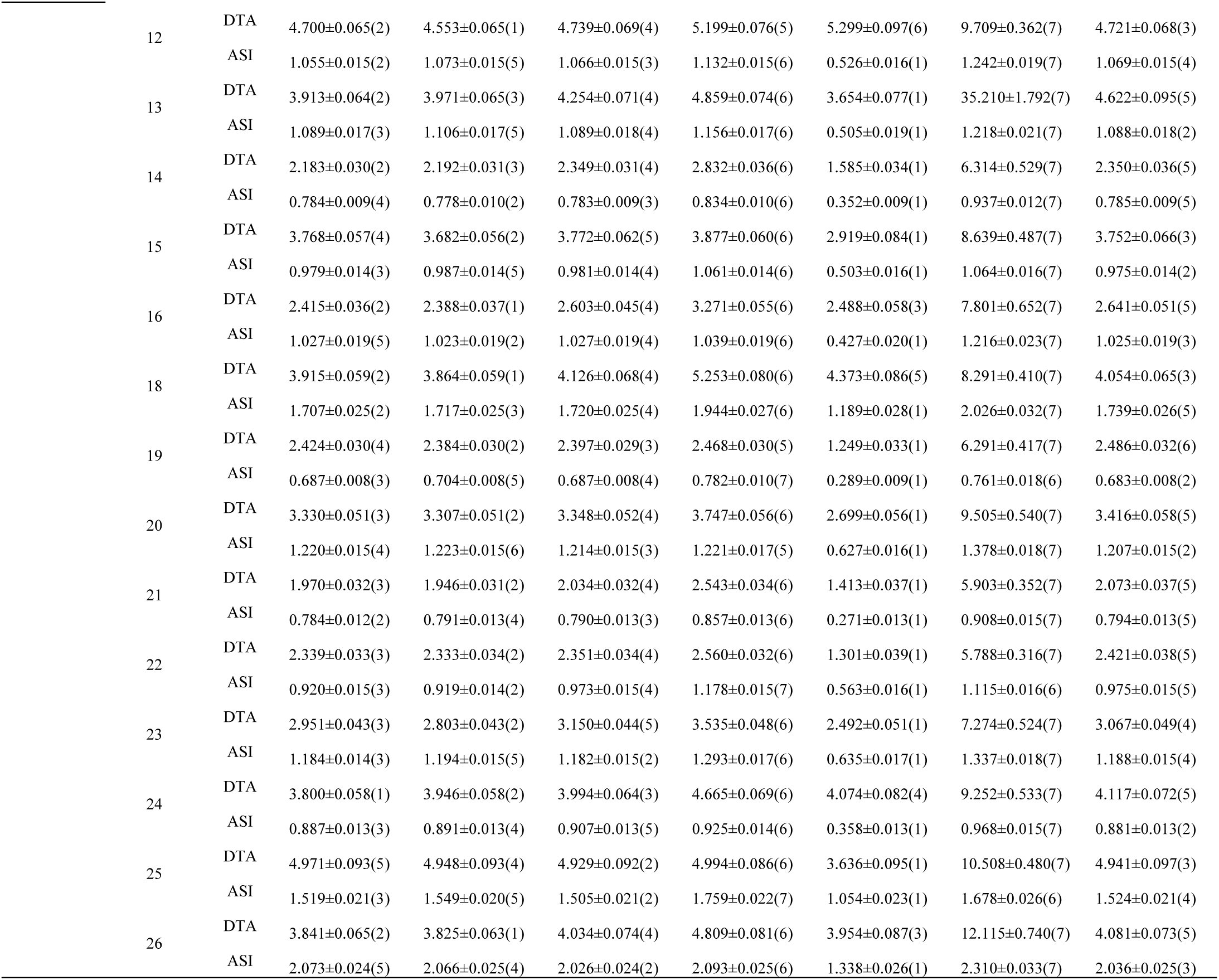
The evaluation of the genomic prediction models (rrBLUP, BayesB, RKHS, RF, SVR, MLP and Ensemble) for the days to anthesis (DTA) and anthesis to silking interval (ASI) traits at the population level in both TeoNAM and MaizeNAM datasets. The prediction performance was measured in a) Pearson correlation and b) mean squared error (MSE). The sign “±” indicates standard error. Values inside brackets represent the ranking of each genomic prediction model under each population-by-trait prediction scenario and the prediction performance improves as the ranking values become smaller.

**Table S3:** Pearson correlation of each pair of individual genomic prediction models at the level of the predicted phenotypes (the top right triangle) and the genomic marker effects (the bottom left triangle) for the days to anthesis (DTA) trait in the a) TeoNAM and b) MaizeNAM datasets. Each value represents the correlation of its corresponding subplot in Figure S7.

|  | rrBLUP | BayesB | RKHS | RF | SVR | MLP |
| --- | --- | --- | --- | --- | --- | --- |
| rrBLUP |  | 0.976 | 0.807 | 0.823 | 0.408 | 0.525 |
| BayesB | 0.738 |  | 0.762 | 0.884 | 0.381 | 0.512 |
| RKHS | 0.396 | 0.497 |  | 0.569 | 0.619 | 0.452 |
| RF | 0.355 | 0.629 | 0.406 |  | 0.310 | 0.408 |
| SVR | 0.143 | 0.258 | 0.559 | 0.332 |  | 0.231 |
| MLP | 0.062 | 0.062 | 0.071 | 0.036 | 0.062 |  |

b)
|  | **rrBLUP** | **BayesB** | **RKHS** | **RF** | **SVR** | **MLP** |
| --- | --- | --- | --- | --- | --- | --- |
| **rrBLUP** |  | 0.996 | 0.992 | 0.958 | 0.944 | 0.807 |
| **BayesB** | 0.866 |  | 0.988 | 0.962 | 0.940 | 0.804 |
| **RKHS** | 0.075 | 0.061 |  | 0.975 | 0.972 | 0.787 |
| **RF** | 0.452 | 0.629 | 0.048 |  | 0.970 | 0.749 |
| **SVR** | 0.508 | 0.541 | 0.066 | 0.647 |  | 0.726 |
| **MLP** | 0.457 | 0.437 | 0.029 | 0.209 | 0.243 |  |

**Table S4:** Pearson correlation of each individual genomic prediction model pair at the level of the predicted phenotypes (the top right triangle) and the genomic marker effects (the bottom left triangle) for the anthesis to silking interval (ASI) trait in the a) TeoNAM and b) MaizeNAM datasets. Each value represents the correlation of its corresponding subplot in Figure S8.

|  | **rrBLUP** | **BayesB** | **RKHS** | **RF** | **SVR** | **MLP** |
| --- | --- | --- | --- | --- | --- | --- |
| **rrBLUP** |  | 0.991 | 0.942 | 0.839 | 0.850 | 0.661 |
| **BayesB** | 0.899 |  | 0.931 | 0.841 | 0.842 | 0.654 |
| **RKHS** | 0.777 | 0.716 |  | 0.927 | 0.896 | 0.715 |
| **RF** | 0.437 | 0.561 | 0.417 |  | 0.845 | 0.718 |
| **SVR** | 0.532 | 0.508 | 0.724 | 0.397 |  | 0.621 |
| **MLP** | 0.356 | 0.341 | 0.353 | 0.193 | 0.267 |  |

b)
|  | **rrBLUP** | **BayesB** | **RKHS** | **RF** | **SVR** | **MLP** |
| --- | --- | --- | --- | --- | --- | --- |
| **rrBLUP** |  | 0.991 | 0.985 | 0.895 | 0.923 | 0.874 |
| **BayesB** | 0.902 |  | 0.979 | 0.913 | 0.921 | 0.867 |
| **RKHS** | 0.062 | 0.055 |  | 0.928 | 0.965 | 0.883 |
| **RF** | 0.499 | 0.643 | 0.047 |  | 0.933 | 0.820 |
| **SVR** | 0.584 | 0.652 | 0.058 | 0.698 |  | 0.845 |
| **MLP** | 0.712 | 0.666 | 0.053 | 0.425 | 0.542 |  |

**Fig. S1:**
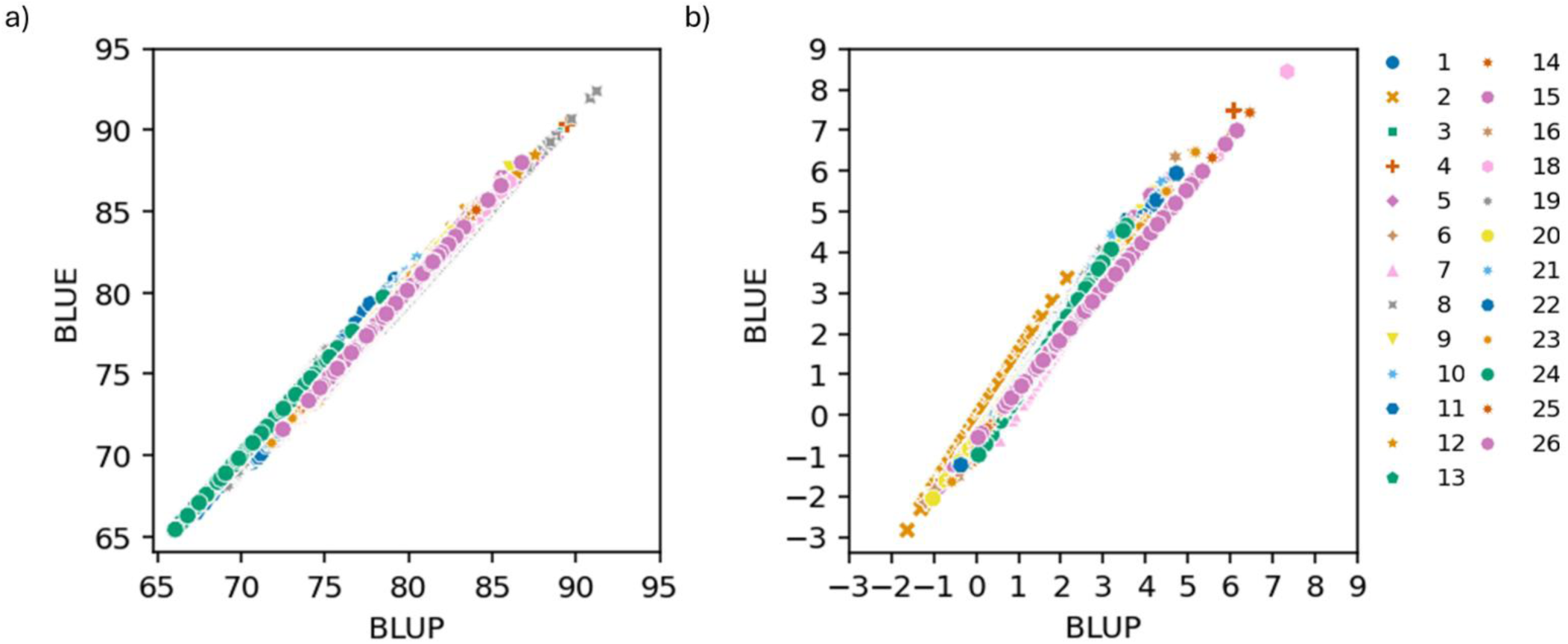
Scatter plots comparing the best linear unbiased predictors (BLUP) and the best linear unbiased estimates (BLUE) at the population level for a) the days to anthesis (DTA) and b) anthesis to silking interval (ASI) traits in the MaizeNAM dataset. The numbers in the legend indicate the population number in the MaizeNAM dataset, showing 25 populations in total.

**Fig. S2:**
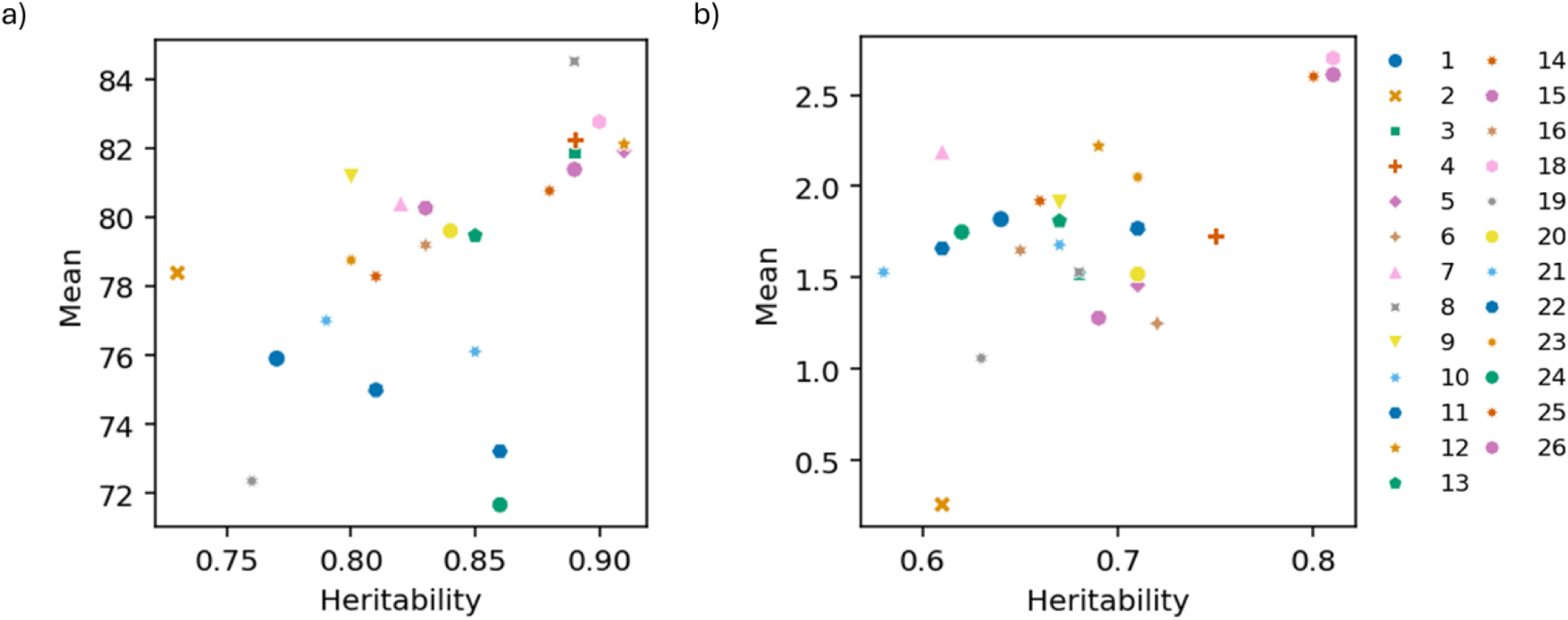
Scatter plots comparing heritability and mean phenotype value in each population for a) the days to anthesis (DTA) and b) anthesis to silking interval (ASI) traits in the MaizeNAM dataset. The numbers in the legend indicate the population number in the MaizeNAM dataset, showing 25 populations in total.

**Fig. S3:**
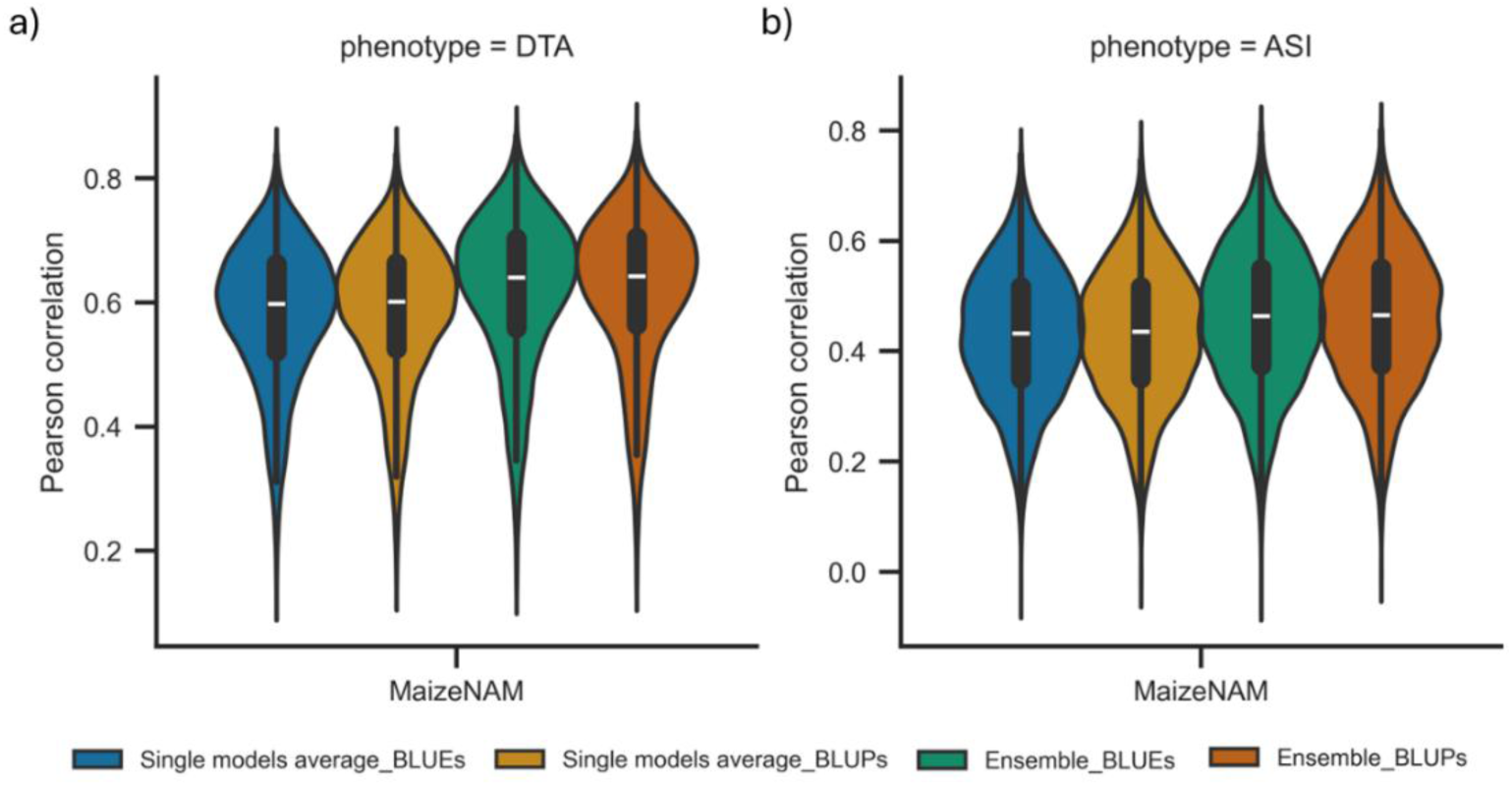
The comparison of Pearson correlation for predicting the best linear unbiased estimates (BLUEs) and the best linear unbiased predictions (BLUPs) between the average of the six individual genomic prediction models (rrBLUP, BayesB, RKHS, RF, SVR and MLP) (Single models average) and the ensemble-average model (Ensemble) in the MaizeNAM dataset. a) the days to anthesis (DTA) and b) anthesis to silking interval (ASI) traits were targeted for the performance comparison. The prediction performance was measured in 3,750 prediction scenarios for each trait. The prediction scenarios were generated by the combination of the three training-test ratios (0.8-0.2, 0.65-0.35 and 0.5-0.5), populations and sampling numbers. The width of the violins represents the distribution of performance metrics. The white horizontal lines on the black box plots show the median value for each metric. The whiskers extend 1.5 times the interquartile range.

**Fig. S4:**
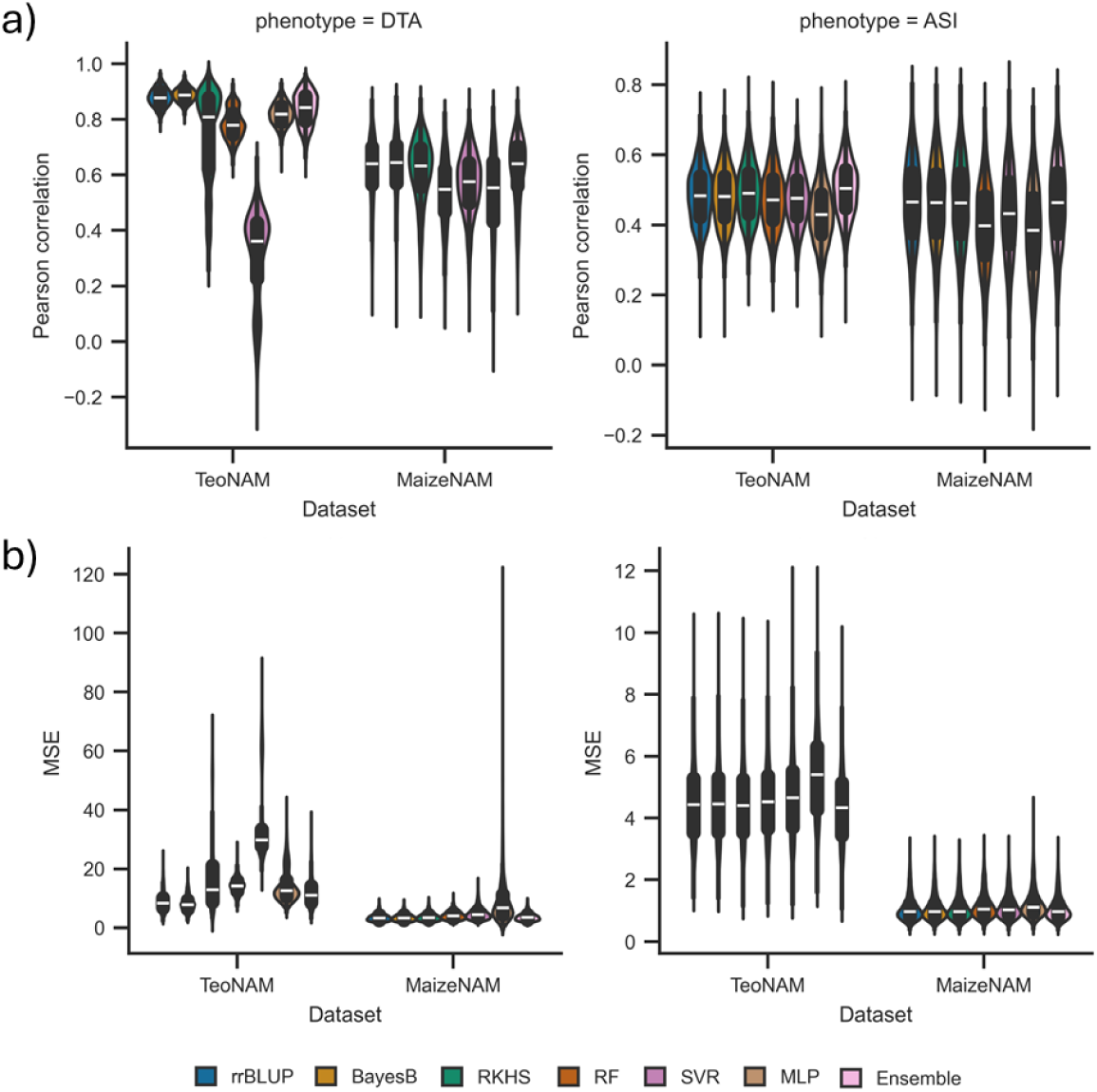
The comparison of prediction performance between the six individual genomic prediction models (rrBLUP, BayesB, RKHS, RF, SVR and MLP) and the ensemble-average model (Ensemble) for the days to anthesis (DTA) and anthesis to silking interval (ASI) traits in the TeoNAM and MaizeNAM datasets. The prediction performance was measured using the two metrics: a) Pearson correlation and b) mean squared error (MSE). The prediction performance was measured in 7,500 prediction scenarios for the TeoNAM dataset and 3,750 prediction scenarios for the MaizeNAM dataset for each trait. The prediction scenarios were generated by the combination of the three training-test ratios (0.8-0.2, 0.65-0.35 and 0.5-0.5), populations and sampling numbers. The width of the violins represents the distribution of performance metrics. The white horizontal lines on the black box plots show the median value for each metric. The whiskers extend 1.5 times the interquartile range.

**Fig. S5:**
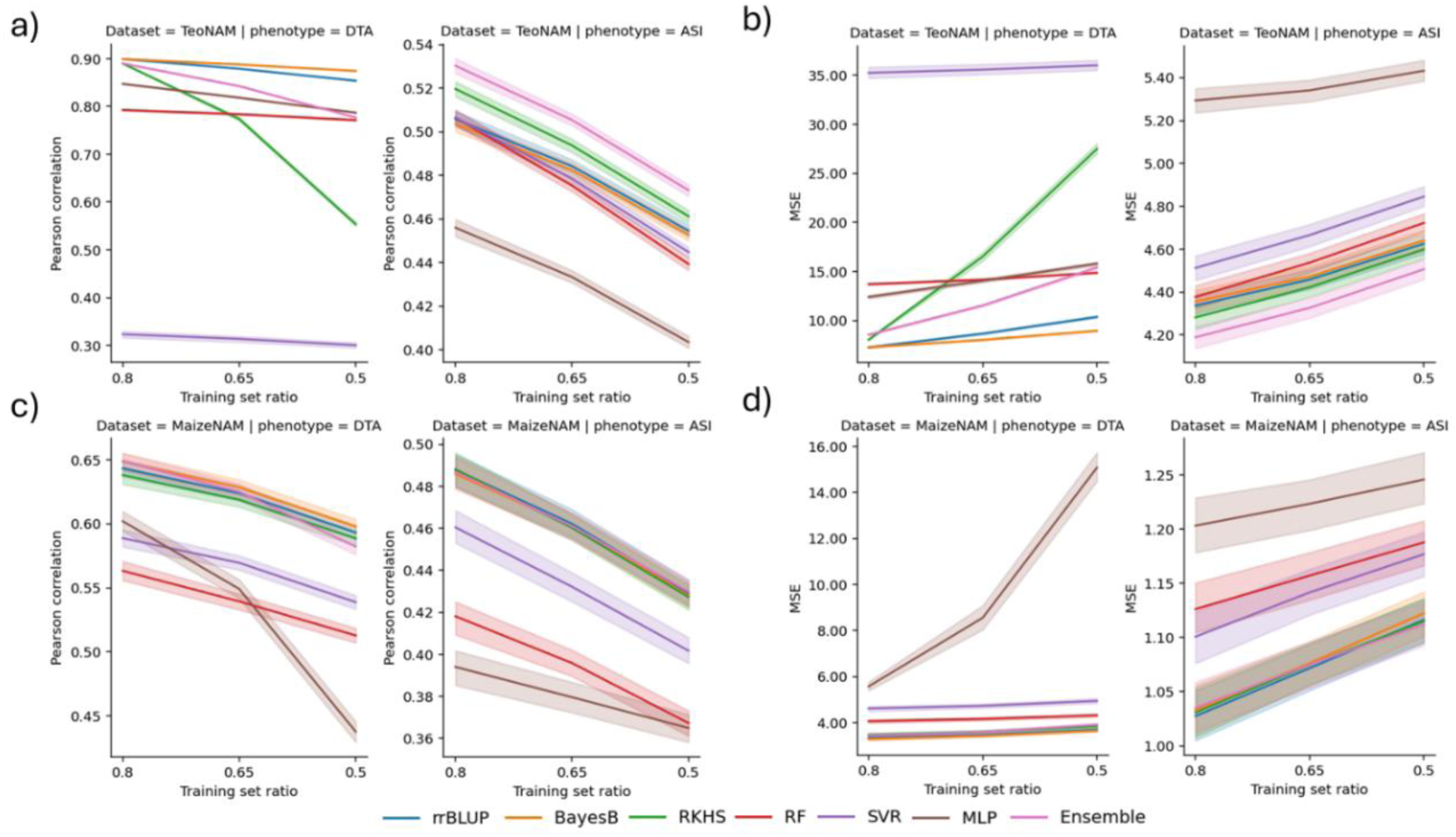
The performance metric transition of the genomic prediction models (rrBLUP, BayesB, RKHS, RF, SVR, MLP and Ensemble) over the ratio reduction of training set for the days to anthesis (DTA) and anthesis to silking interval (ASI) traits: a) Pearson correlation and b) mean squared error (MSE) in the TeoNAM dataset and c) Pearson correlation and d) MSE in the MaizeNAM dataset. In each subplot, the x-axis represents the ratio of the training set whereas the y-axis indicates corresponding metric values. Intervals surrounding each line illustrates 95% the confidence interval.

**Fig. S6:**
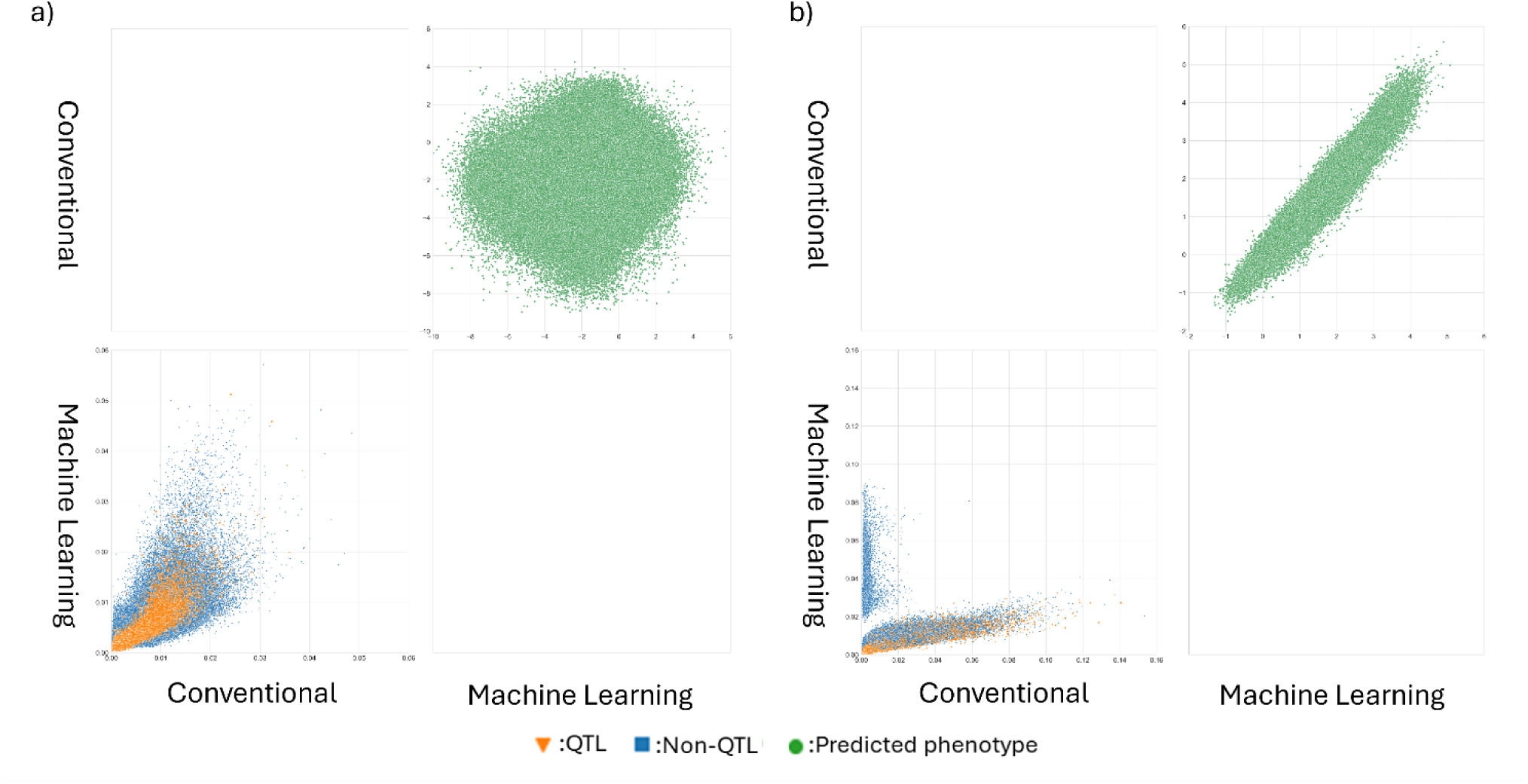
Pairwise comparisons of the conventional (rrBLUP, BayesB and RKHS) and machine learning models (RF, SVR and MLP) for the anthesis to silking interval (ASI) trait across all the prediction scenarios for the a) TeoNAM (7,500 prediction scenarios) and b) MaizeNAM (3,750 prediction scenarios) datasets. The prediction scenarios were generated by the combination of the three training-test ratios (0.8-0.2, 0.65-0.35 and 0.5-0.5), populations and sampling numbers. The genomic prediction model groups were compared for mean predicted phenotypes (top right triangle) and mean normalised genomic marker effects (the bottom left triangle) calculated within each prediction model category. The green dots represent a pair of predicted phenotypes of RIL samples in the test sets for each prediction scenario. The blue squares and orange triangles represent a pair of extracted genomic marker effects from each genomic marker in each sample scenario that were identified as non-QTL and QTL markers, respectively, by Chen et al. (2019) for the TeoNAM dataset and Buckler et al. (2009) for the MaizeNAM dataset.

**Fig. S7:**
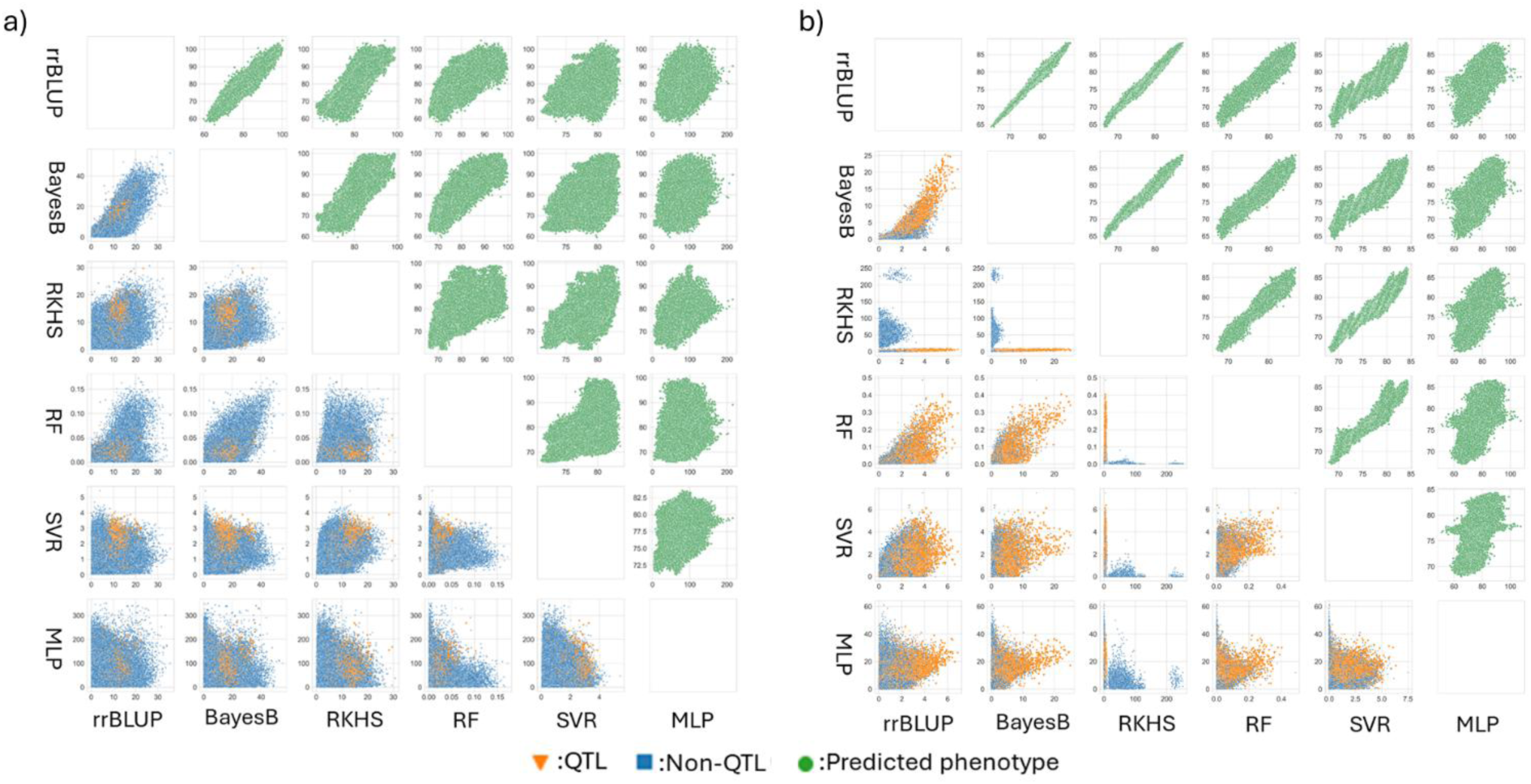
Pairwise comparisons of the individual genomic prediction models for the days to anthesis (DTA) trait across all the prediction scenarios for the a) TeoNAM (7,500 prediction scenarios) and b) MaizeNAM (3,750 prediction scenarios) datasets. The prediction scenarios were generated by the combination of the three training-test ratios (0.8-0.2, 0.65-0.35 and 0.5-0.5), populations and sampling numbers. The genomic prediction models were compared at the level of predicted phenotypes (top right triangle) and genomic marker effects (the bottom left triangle). The green dots represent a pair of predicted phenotypes of RIL samples in the test sets for each prediction scenario. The blue squares and orange triangles represent a pair of extracted genomic marker effects from each genomic marker in each sample scenario that were identified as non-QTL and QTL markers, respectively, by Chen et al. (2019) for the TeoNAM dataset and Buckler et al. (2009) for the MaizeNAM dataset.

**Fig. S8:**
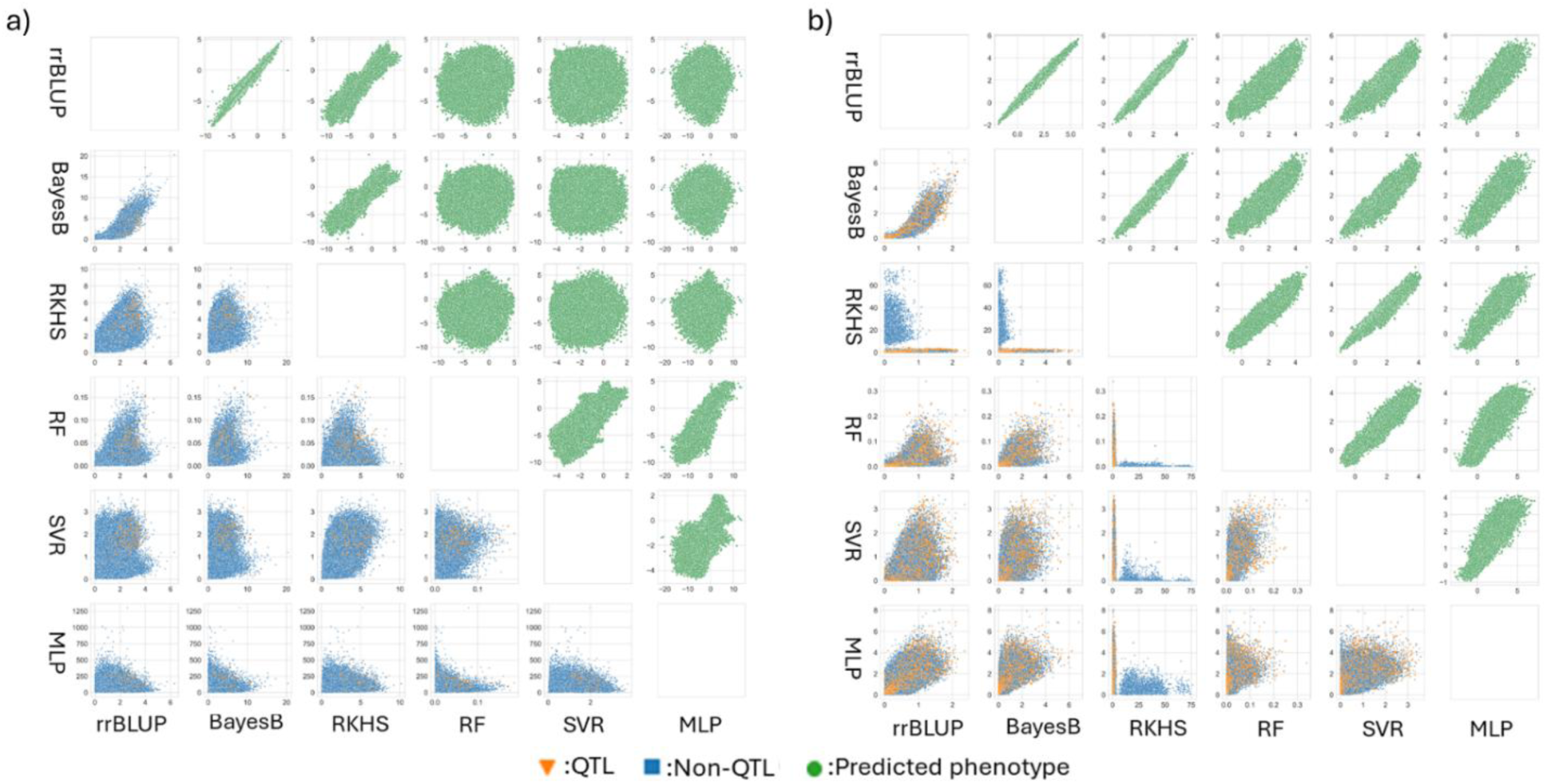
Pairwise comparisons of the individual genomic prediction models for the anthesis to silking interval (ASI) trait across all the prediction scenarios for the a) TeoNAM (7,500 prediction scenarios) and b) MaizeNAM (3,750 prediction scenarios) datasets. The prediction scenarios were generated by the combination of the three training-test ratios (0.8-0.2, 0.65-0.35 and 0.5-0.5), populations and sampling numbers. The genomic prediction models were compared at the level of predicted phenotypes (top right triangle) and genomic marker effects (the bottom left triangle). The green dots represent a pair of predicted phenotypes of RIL samples in the test sets for each prediction scenario. The blue squares and orange triangles represent a pair of extracted genomic marker effects from each genomic marker in each sample scenario that were identified as non-QTL and QTL markers, respectively, by Chen et al. (2019) for the TeoNAM dataset and Buckler et al. (2009) for the MaizeNAM dataset.

**Fig. S9:**
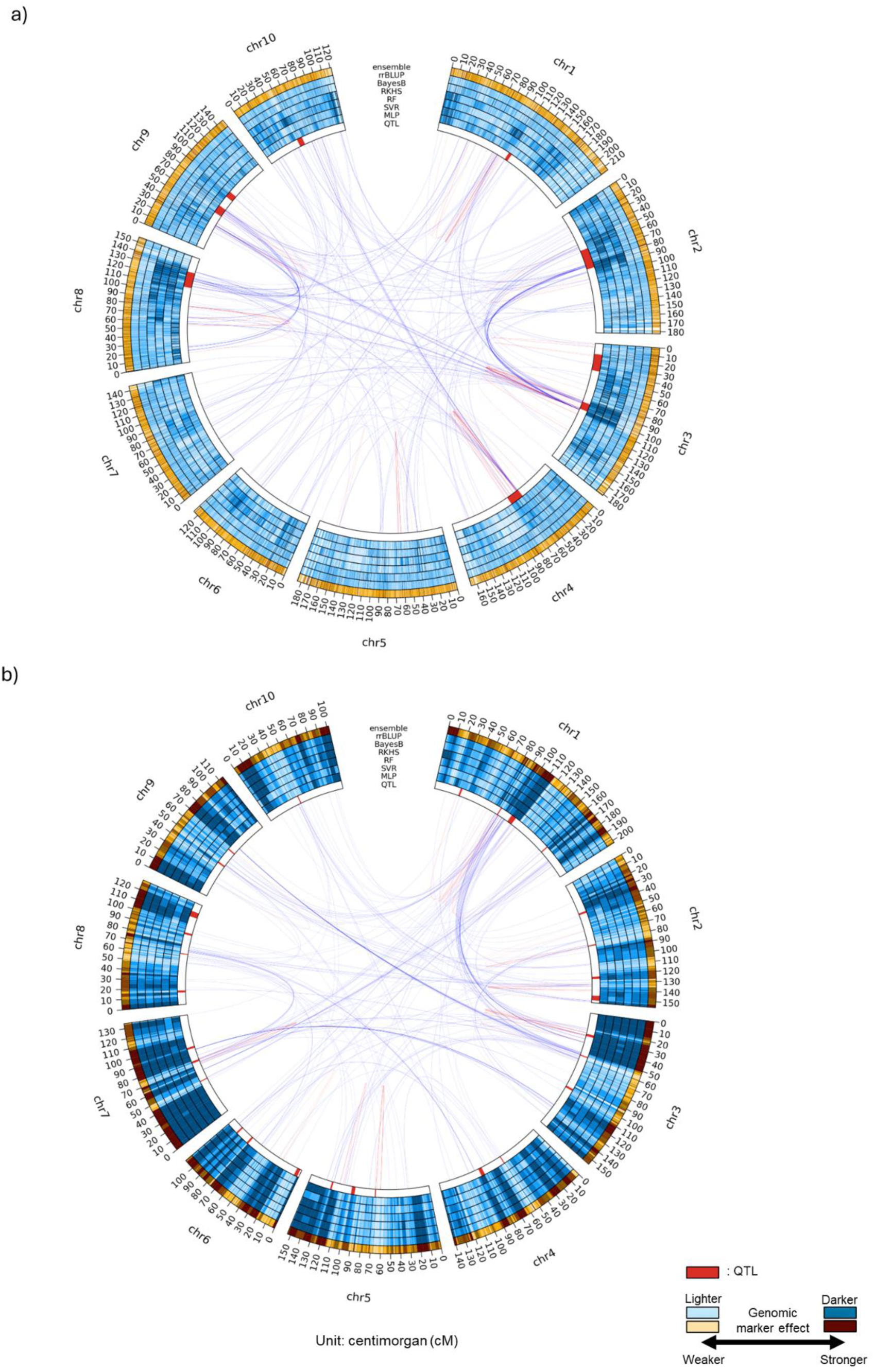
Circos plots for the anthesis to silking interval (ASI) trait for the a) TeoNAM and b) MaizeNAM datasets. The innermost (QTL) ring shows the QTL gene regions estimated by Chen et al. (2019) for the TeoNAM dataset and by Buckler et al. (2009) for the MaizeNAM dataset. The blue rings (second to seventh) represent the genomic marker effects across the gene regions estimated by MLP, SVR, RF, RKHS, BayesB and rrBLUP, respectively. The outermost orange ring is the genomic marker effects for the ensemble-average model (ensemble). The numbers at the outermost ring represent genetic distance in centimorgans (cM). The darkness level of the blue and orange colours indicates the strength of the genomic marker effects, sectioned into ten levels using the quantiles. Darker colours represent higher genomic marker effect levels. The red and blue lines between genome regions are the genomic marker interaction effects calculated by pairwise Shapley scores from RF (top 0.01%; red = within chromosome and blue = between chromosomes).

**Fig. S10:**
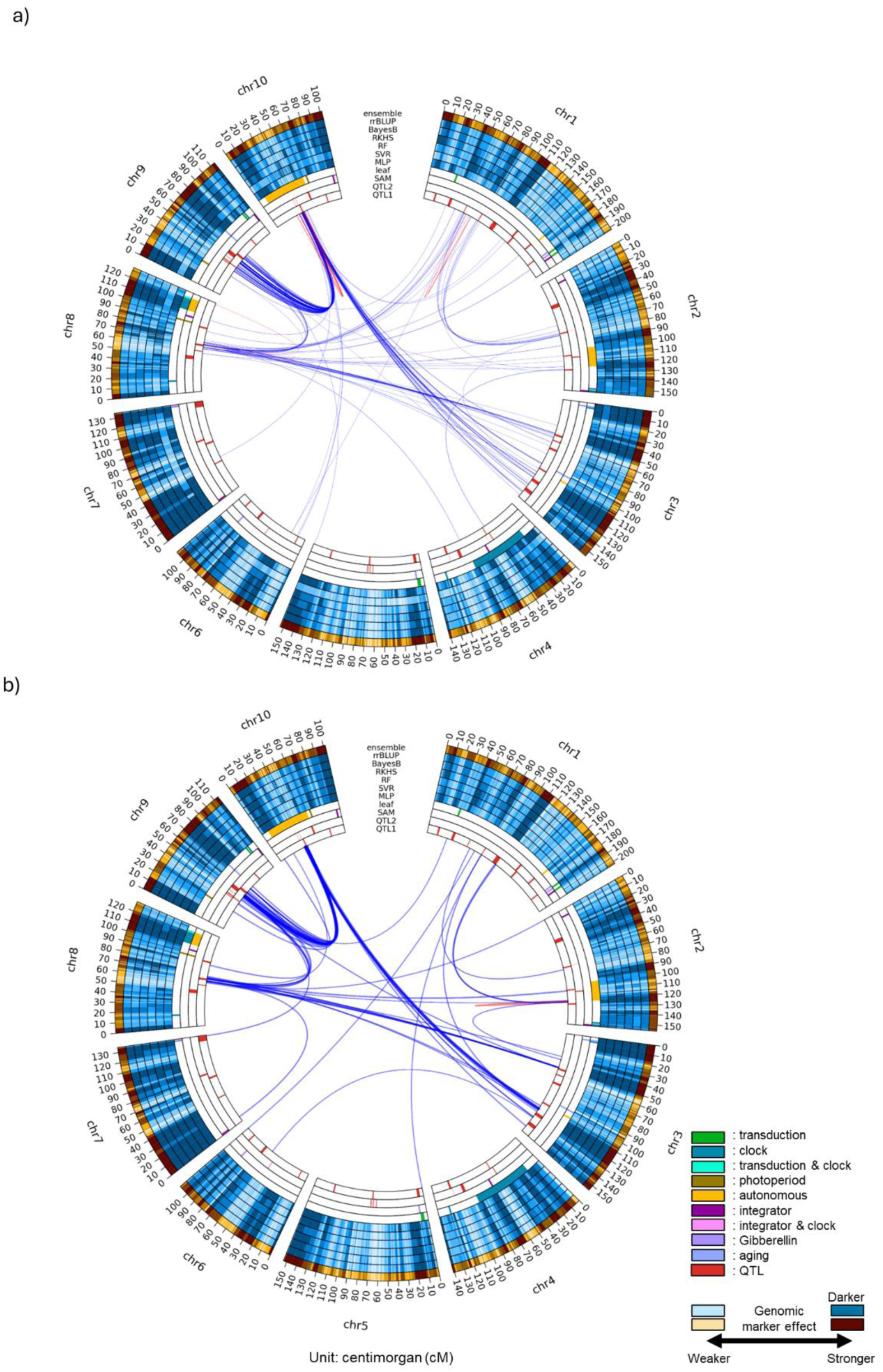
Circos plots for the days to anthesis (DTA) trait for predicting the a) best linear unbiased estimates (BLUEs) and b) best linear unbiased predictions (BLUPs) in MaizeNAM datasets. The innermost (QTL 1) ring shows the QTL gene regions estimated by Chen et al. (2019) for the TeoNAM dataset and by Buckler et al. (2009) for the MaizeNAM dataset. The second innermost ring (QTL 2) represents the QTL gene regions identified by Wisser et al. (2019). The third and fourth innermost rings represent gene regulators that affect the shoot apical meristem (SAM) and leaf, respectively, identified by Dong et al. (2012). The blue rings (fifth to tenth) represent the genomic marker effects across the gene regions estimated by MLP, SVR, RF, RKHS, BayesB and rrBLUP, respectively. The outermost orange ring is the genomic marker effects for the ensemble-average model (ensemble). The numbers at the outermost ring represent genetic distance in centimorgans (cM). The darkness level of the blue and orange colours indicates the strength of the genomic marker effects, sectioned into ten levels using the quantiles. Darker colours represent higher genomic marker effect levels. The red and blue lines between genome regions are the genomic marker interaction effects calculated by pairwise Shapley scores from RF (top 0.01%; red = within chromosome and blue = between chromosomes).

**Fig. S11:**
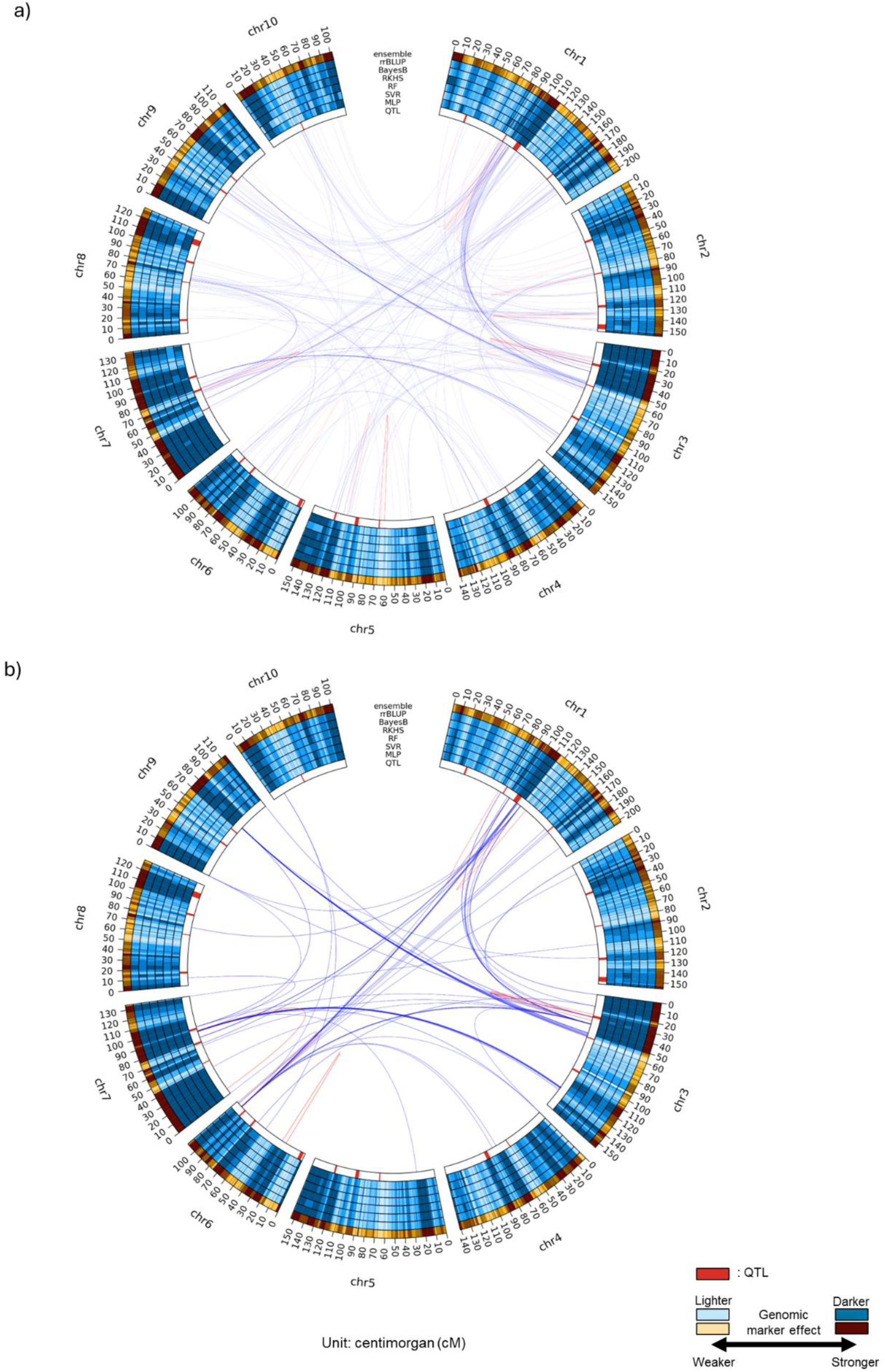
Circos plots for the anthesis to silking interval (ASI) trait for predicting the a) best linear unbiased estimates (BLUEs) and b) best linear unbiased predictions (BLUPs) in MaizeNAM datasets. The innermost (QTL) ring shows the QTL gene regions estimated by Chen et al. (2019) for the TeoNAM dataset and by Buckler et al. (2009) for the MaizeNAM dataset. The blue rings (second to seventh) represent the genomic marker effects across the gene regions estimated by MLP, SVR, RF, RKHS, BayesB and rrBLUP, respectively. The outermost orange ring is the genomic marker effects for the ensemble-average model (ensemble). The numbers at the outermost ring represent genetic distance in centimorgans (cM). The darkness level of the blue and orange colours indicates the strength of the genomic marker effects, sectioned into ten levels using the quantiles. Darker colours represent higher genomic marker effect levels. The red and blue lines between genome regions are the genomic marker interaction effects calculated by pairwise Shapley scores from RF (top 0.01%; red = within chromosome and blue = between chromosomes).

## References

1. Alemu A, Åstrand J, Montesinos-Lopez OA, y Sanchez JI, Fernandez-Gonzalez J, Tadesse W, Vetukuri RR, Carlsson AS, Ceplitis A, Crossa J et al. 2024. Genomic selection in plant breeding: Key factors shaping two decades of progress. Molecular Plant. 17:552–578. 10.1016/j.molp.2024.03.007.

2. Azodi CB, Bolger E, McCarren A, Roantree M, de Los Campos G, Shiu SH. 2019. Benchmarking parametric and machine learning models for genomic prediction of complex traits. G3: Genes, Genomes, Genetics. 9:3691–3702. 10.1534/g3.119.400498.

3. Bernardo R, Yu J. 2007. Prospects for genomewide selection for quantitative traits in maize. Crop Science. 47:1082–1090. 10.2135/cropsci2006.11.0690.

4. Bian Y, Chen H. 2021. When does diversity help generalization in classification ensembles? IEEE Transactions on Cybernetics. 52:9059–9075. 10.1109/TCYB.2021.3053165.

5. Breiman L. 2001. Random forests. Machine Learning. 45:5–32. 10.1023/A:1010933404324.

6. Buckler ES, Holland JB, Bradbury PJ, Acharya CB, Brown PJ, Browne C, Ersoz E, Flint-Garcia S, Garcia A, Glaubitz JC et al. 2009. The genetic architecture of maize flowering time. Science. 325:714–718.

7. Butler D, Cullis B, Gilmour A, Gogel B, Thompson R. 2017. Asreml-r reference manual version 4. VSN International Ltd, Hemel Hempstead, HP1 1ES, UK.

8. Ceccarelli S, Grando S, Maatougui M, Michael M, Slash M, Haghparast R, Rahmanian M, Taheri A, Al-Yassin A, Benbelkacem A et al. 2010. Plant breeding and climate changes. The Journal of Agricultural Science. 148:627–637. 10.1017/S0021859610000651.

9. Chang CC, Chow CC, Tellier LC, Vattikuti S, Purcell SM, Lee JJ. 2015. Second-generation plink: rising to the challenge of larger and richer datasets. GigaScience. 4. 10.1186/s13742015-0047-8.

10. Chen Q, Yang CJ, York AM, Xue W, Daskalska LL, DeValk CA, Krueger KW, Lawton SB, Spiegelberg BG, Schnell JM et al. 2019. Teonam: A nested association mapping population for domestication and agronomic trait analysis in maize. Genetics. 213:1065–1078. 10.1534/genetics.119.302594.

11. Cooper M, Gho C, Leafgren R, Tang T, Messina C 2014b. Breeding drought-tolerant maize hybrids for the US corn-belt: discovery to product. Journal of Experimental Botany 65:6191–6204. 10.1093/jxb/eru064.

12. Cooper M, Messina CD. 2023. Breeding crops for drought-affected environments and improved climate resilience. The Plant Cell. 35:162–186. 10.1093/plcell/koac321.

13. Cooper M, Messina CD, Podlich D, Totir LR, Baumgarten A, Hausmann NJ, Wright D, Graham G 2014a. Predicting the future of plant breeding: complementing empirical evaluation with genetic prediction. Crop and Pasture Science 65:311–336. 10.1071/CP14007.

14. Cooper M, Tomura S, Wilkinson MJ, Powell O, Messina CD. 2025. Breeding perspectives on tackling trait genome-to-phenome (g2p) dimensionality using ensemble-based genomic prediction. Theoretical and Applied Genetics. 10.1007/s00122-025-04960-6.

15. Crossa J, Montesinos-Lopez OA, Costa-Neto G, Vitale P, Martini JW, Runcie D, Fritsche-Neto R, Montesinos-Lopez A, Pérez-Rodríguez P, Gerard G et al. 2025. Machine learning algorithms translate big data into predictive breeding accuracy. Trends in Plant Science. 10.1016/j.tplants.2024.09.011.

16. Dave VS, Dutta K. 2014. Neural network based models for software effort estimation: a review. Artificial Intelligence Review. 42:295–307. 10.1007/s10462-012-9339-x.

17. de los Campos G, Gianola D, Rosa GJ. 2009. Reproducing kernel hilbert spaces regression: a general framework for genetic evaluation. Journal of animal science. 87:1883–1887. 10.2527/jas.2008-1259.

18. Diepenbrock C, Tang T, Jines M, Technow F, Lira S, Podlich D, Cooper M, Messina C (2022) Can we harness digital technologies and physiology to hasten genetic gain in U.S. maize breeding? Plant Physiology 188:1141–1157. 10.1093/plphys/kiab527.

19. Dong Z, Danilevskaya O, Abadie T, Messina C, Coles N, Cooper M. 2012. A gene regulatory network model for floral transition of the shoot apex in maize and its dynamic modeling. Plos One. 10.1371/journal.pone.0043450.

20. Drucker H, Burges CJ, Kaufman L, Smola A, Vapnik V. 1996. Support vector regression machines. Advances in neural information processing systems. 9.

21. Endelman JB. 2011. Ridge regression and other kernels for genomic selection with r package rrblup. The Plant Genome. 4:250–255. 10.3835/plantgenome2011.08.0024.

22. Escamilla DM, Li D, Negus KL, Kappelmann KL, Kusmec A, Vanous AE, Schnable PS, Li X, Yu J. 2025. Genomic selection: Essence, applications, and prospects. The Plant Genome 18:e70053. 10.1002/tpg2.70053

23. Gianola D, Van Kaam JB. 2008. Reproducing kernel hilbert spaces regression methods for genomic assisted prediction of quantitative traits. Genetics. 178:2289–2303. 10.1534/genetics.107.084285.

24. Gibbs PM, Paril JF, Fournier-Level A. 2025. Trait genetic architecture and population structure determine model selection for genomic prediction in natural arabidopsis thaliana populations. Genetics. 229:iyaf003. 10.1093/genetics/iyaf003.

25. Hammer G, Cooper M, Tardieu F, Welch S, Walsh B, van Eeuwijk F, Chapman S, Podlich D. 2006. Models for navigating biological complexity in breeding improved crop plants. Trends in plant science. 11:587–593. 10.1016/j.tplants.2006.10.006.

26. Heffner EL, Lorenz AJ, Jannink JL, Sorrells ME. 2010. Plant breeding with genomic selection: gain per unit time and cost. Crop science. 50:1681–1690. 10.2135/cropsci2009.11.0662.

27. Heilmann PG, Frisch M, Abbadi A, Kox T, Herzog E. 2023. Stacked ensembles on basis of parentage information can predict hybrid performance with an accuracy comparable to marker-based gblup. Frontiers in Plant Science. 14:1178902.

28. Heslot N, Yang HP, Sorrells ME, Jannink JL. 2012. Genomic selection in plant breeding: a comparison of models. Crop science. 52:146–160. 10.2135/cropsci2011.06.0297.

29. Hong, L, Page, SE. 2004. Groups of diverse problem solvers can outperform groups of high-ability problem solvers. Proceedings of the National Academy of Sciences, 101(46), 16385–16389. 10.1073/pnas.040372310.

30. Howard R, Lipka AE. 2025. Genomic selection and reproducibility: are complex models distracting us from true scientific validity in the presence of genotype-by-environment interaction? G3: Genes, Genomes, Genetics. 15:jkaf244. 10.1093/g3journal/jkaf244.

31. Huang C, Sun H, Xu D, Chen Q, Liang Y, Wang X, Xu G, Tian J, Wang C, Li D et al. (2018) Zmcct9 enhances maize adaptation to higher latitudes. Proceedings of the National Academy of Sciences. 115:E334–E341. 10.1073/pnas.1718058115.

32. Hufford MB, Xu X, Van Heerwaarden J, Pyhäjärvi T, Chia JM, Cartwright RA, Elshire RJ, Glaubitz JC, Guill KE, Kaeppler SM et al. 2012. Comparative population genomics of maize domestication and improvement. Nature genetics. 44:808–811. 10.1038/ng.2309.

33. Ishwaran H. 2015. The effect of splitting on random forests. Machine learning. 99:75–118. 10.1007/s10994-014-5451-2.

34. Jeon D, Kang Y, Lee S, Choi S, Sung Y, Lee TH, Kim C. 2023. Digitalizing breeding in plants: A new trend of next-generation breeding based on genomic prediction. Frontiers in Plant Science. 14:1092584. 10.3389/fpls.2023.1092584.

35. John M, Haselbeck F, Dass R, Malisi C, Ricca P, Dreischer C, Schultheiss SJ, Grimm DG. 2022. A comparison of classical and machine learning-based phenotype prediction methods on simulated data and three plant species. Frontiers in Plant Science. 13:932512. 10.3389/fpls.2022.932512.

36. Khaipho-Burch M, Cooper M, Crossa J, de Leon N, Holland J, Lewis R, McCouch S, Murray SC, Rabbi I, Ronald P, Ross-Ibarra J, Weigel D, Buckler ES (2023) Scale up trials to validate modified crops’ benefits. Nature 621:470–473. 10.1038/d41586-023-02895-w.

37. Kholová J, Urban MO, Cock J, Arcos J, Arnaud E, Aytekin D, Azevedo V, Barnes AP, Ceccarelli S, Chavarriaga P et al. 2021. In pursuit of a better world: crop improvement and the cgiar. Journal of Experimental Botany. 72:5158–5179. 10.1093/jxb/erab226.

38. Kick DR, Washburn JD. 2023. Ensemble of best linear unbiased predictor, machine learning, and deep learning models predict maize yield better than each model alone. in silico Plants. p. diad015. 10.1093/insilicoplants/diad015.

39. Krzywinski M, Schein J, Birol I, Connors J, Gascoyne R, Horsman D, Jones SJ, Marra MA. 2009. Circos: an information aesthetic for comparative genomics. Genome research. 19:1639–1645. 10.1126/science.1174276.

40. Langridge P, Braun H, Hulke B, Ober E, Prasanna B. 2021. Breeding crops for climate resilience. Theoretical and Applied Genetics. 134:1607–1611. 10.1007/s00122-021-03854-7.

41. Loshchilov I, Hutter F. 2017. Decoupled weight decay regularization. arXiv preprint arXiv:1711.05101. 10.48550/arXiv.1711.05101.

42. Lourenço VM, Ogutu JO, Rodrigues RA, Posekany A, Piepho HP. 2024. Genomic prediction using machine learning: a comparison of the performance of regularized regression, ensemble, instance-based and deep learning methods on synthetic and empirical data. BMC genomics. 25:152. 10.1186/s12864-023-09933-x.

43. Lundberg SM, Lee SI. 2017. A unified approach to interpreting model predictions. Advances in neural information processing systems. 30.

44. Meher PK, Rustgi S, Kumar A. 2022. Performance of bayesian and blup alphabets for genomic prediction: analysis, comparison and results. Heredity. 128:519–530. 10.1038/s41437-02200539-9.

45. Messina C, Garcia-Abadillo J, Powell O, Tomura S, Zare A, Ganapathysubramanian B, Cooper M. 2025. Toward a general framework for ai-enabled prediction in crop improvement. Theoretical and Applied Genetics. 138:1–15. 10.1007/s00122-025-04928-6.

46. Messina CD, Gho C, Hammer GL, Tang T, Cooper M. 2023. Two decades of harnessing standing genetic variation for physiological traits to improve drought tolerance in maize. Journal of experimental botany. 74:4847–4861. 10.1093/jxb/erad231.

47. Messina CD, Technow F, Tang T, Totir R, Gho C, Cooper M. 2018. Leveraging biological insight and environmental variation to improve phenotypic prediction: Integrating crop growth models (cgm) with whole genome prediction (wgp). European Journal of Agronomy. 100:151–162. 10.1016/j.eja.2018.01.007.

48. Meuwissen TH, Hayes BJ, Goddard M. 2001. Prediction of total genetic value using genome-wide dense marker maps. genetics. 157:1819–1829. 10.1093/genetics/157.4.1819.

49. Molnar C. 2020. Interpretable machine learning. Lulu. com.

50. Montesinos-Lopez A, Crespo-Herrera L, Dreisigacker S, Gerard G, Vitale P, Saint Pierre C, Govindan V, Tarekegn ZT, Flores MC, Pérez-Rodríguez P et al. 2024. Deep learning methods improve genomic prediction of wheat breeding. Frontiers in Plant Science. 15:1324090. 10.3389/fpls.2024.1324090.

51. Montesinos-López A, Montesinos-López OA, Ramos-Pulido S, Mosqueda-González BA, GuerreroArroyo EA, Crossa J, Ortiz R. 2025. Artificial intelligence meets genomic selection: comparing deep learning and gblup across diverse plant datasets. Frontiers in Genetics. 16:1568705. 10.3389/fgene.2025.1568705.

52. Nair V, Hinton GE. 2010. Rectified linear units improve restricted boltzmann machines. pp. 807–814.

53. Nascimento M, Nascimento ACC, Azevedo CF, Oliveira ACBd, Caixeta ET, Jarquin D. 2024. Enhancing genomic prediction with stacking ensemble learning in arabica coffee. Frontiers in Plant Science. 15:1373318. 10.3389/fpls.2024.1373318.

54. Page SE. 2007. Making the difference: Applying a logic of diversity. Academy of Management Perspectives 21.4: 6–20. 10.5465/amp.2007.27895335.

55. Page SE. 2018. The model thinker: What you need to know to make data work for you. Hachette UK.

56. Pérez P, de Los Campos G. 2014. Genome-wide regression and prediction with the bglr statistical package. Genetics. 198:483–495. 10.1534/genetics.114.164442.

57. Pixley KV, Cairns JE, Lopez-Ridaura S, Ojiewo CO, Dawud MA, Drabo I, Mindaye T, Nebie B, Asea G, Das B et al. 2023. Redesigning crop varieties to win the race between climate change and food security. Molecular Plant. 16:1590–1611. 10.1016/j.molp.2023.09.003.

58. Plavšin I, Gunjača J, Galić V, Novoselović D. 2022. Evaluation of genomic selection methods for wheat quality traits in biparental populations indicates inclination towards parsimonious solutions. Agronomy. 12. 10.3390/agronomy12051126.

59. Rosenblatt F. 1958. The perceptron: a probabilistic model for information storage and organization in the brain. Psychological review. 65:386. 10.1037/h0042519.

60. Schapire RE, Rochery M, Rahim M, Gupta N. 2002. Incorporating prior knowledge into boosting. In: . volume 2. pp. 538–545.

61. Shapley LS. 1953. Stochastic games. Proceedings of the national academy of sciences. 39:1095–1100. 10.1073/pnas.39.10.1095.

62. Silva PC, Sanchez AC, Opazo MA, Mardones LA, Acevedo EA. 2022. Grain yield, anthesis-silking interval, and phenotypic plasticity in response to changing environments: Evaluation in temperate maize hybrids. Field Crops Research. 285:108583. 10.1016/j.fcr.2022.108583.

63. Technow F, Messina CD, Totir LR, Cooper M. 2015. Integrating crop growth models with whole genome prediction through approximate bayesian computation. PloS one. 10:e0130855. 10.1371/journal.pone.0130855.

64. Tieleman T. 2012. Lecture 6.5-rmsprop: Divide the gradient by a running average of its recent magnitude. COURSERA: Neural networks for machine learning. 4:26.

65. Tomura S, Wilkinson MJ, Powell O, Cooper M. 2025a. Ensemble AnalySis with Interpretable Genomic Prediction (EasiGP): Computational Tool for Interpreting Ensembles of Genomic Prediction Models. The Plant Genome. 10.1002/tpg2.70138.

66. Tomura S, Wilkinson MJ, Cooper M, Powell O. 2025b. Improved genomic prediction performance with ensembles of diverse models. G3: Genes, Genomes, Genetics. p. jkaf048. 10.1093/g3journal/jkaf048.

67. Tusell, L, Pérez-Rodríguez, P, Forni, S, Gianola, D. 2014. Model averaging for genome-enabled prediction with reproducing kernel Hilbert spaces: a case study with pig litter size and wheat yield. Journal of animal breeding and genetics, 131(2), 105–115. 10.1111/jbg.12070.

68. Von Rueden L, Mayer S, Beckh K, Georgiev B, Giesselbach S, Heese R, Kirsch B, Pfrommer J, Pick A, Ramamurthy R et al. 2021. Informed machine learning–a taxonomy and survey of integrating prior knowledge into learning systems. IEEE Transactions on Knowledge and Data Engineering. 35:614–633. 10.1109/TKDE.2021.3079836.

69. Voss-Fels KP, Cooper M, Hayes BJ. 2019. Accelerating crop genetic gains with genomic selection. Theoretical and Applied Genetics. 132:669–686. 10.1007/s00122-018-3270-8.

70. Wallach D, Martre P, Liu B, Asseng S, Ewert F, Thorburn PJ, van Ittersum M, Aggarwal PK, Ahmed M, Basso B et al. 2018. Multimodel ensembles improve predictions of crop–environment–management interactions. Global Change Biology. 24. 10.1111/gcb.14411.

71. Washburn JD, Varela JI, Xavier A, Chen Q, Ertl D, Gage JL, Holland JB, Lima DC, Romay MC, Lopez-Cruz M et al. 2025. Global genotype by environment prediction competition reveals that diverse modeling strategies can deliver satisfactory maize yield estimates. Genetics. 229:iyae195. 10.1093/genetics/iyae195.

72. Werner CR, Zaman-Allah M, Assefa T, Cairns JE, Atlin GN. 2025. Accelerating genetic gain through early-stage on-farm sparse testing. Trends in Plant Science. 10.1016/j.tplants.2024.10.010.

73. Wisser RJ, Fang Z, Holland JB, Teixeira JE, Dougherty J, Weldekidan T, de Leon N, Flint-Garcia S, Lauter N, Murray SC et al. 2019. The genomic basis for short-term evolution of environmental adaptation in maize. Genetics. 213:1479–1494. 10.1534/genetics.119.302780.

74. Wolpert DH, Macready WG. 1997. No free lunch theorems for optimization. IEEE transactions on evolutionary computation. 1:67–82. 10.1109/4235.585893.

75. Wood D, Mu T, Webb AM, Reeve HW, Lujan M, Brown G. 2023. A unified theory of diversity in ensemble learning. Journal of machine learning research. 24:1–49.

76. Yadav S, Ross EM, Wei X, Powell O, Hivert V, Hickey LT, Atkin F, Deomano E, Aitken KS, Voss-Fels KP et al. 2023. Optimising clonal performance in sugarcane: leveraging non-additive effects via mate-allocation strategies. Frontiers in Plant Science. 14:1260517. 10.3389/fpls.2023.1260517.

77. Yadav S, Wei X, Joyce P, Atkin F, Deomano E, Sun Y, Nguyen LT, Ross EM, Cavallaro T, Aitken KS et al. 2021. Improved genomic prediction of clonal performance in sugarcane by exploiting non-additive genetic effects. Theoretical and Applied Genetics. 134:2235–2252. 10.1007/s00122-021-03822-1.

78. Yang J, Benyamin B, McEvoy BP, Gordon S, Henders AK, Nyholt DR, Madden PA, Heath AC, Martin NG, Montgomery GW et al. 2010. Common snps explain a large proportion of the heritability for human height. Nature genetics. 42:565–569. 10.1038/ng.608.

